# Self-generated chemotaxis of mixed cell populations

**DOI:** 10.1101/2024.12.19.628881

**Authors:** Mehmet Can Uçar, Zane Alsberga, Jonna Alanko, Michael Sixt, Edouard Hannezo

**Author notes:** These authors jointly supervised this work (Michael Sixt, Edouard Hannezo).

## Abstract

Cell and tissue movement in development, cancer invasion, and immune response relies on chemical or mechanical guidance cues. In many systems, this behavior is locally guided by self-generated signaling gradients rather than long-range, pre-patterned cues. However, how heterogeneous mixtures of cells interact non-reciprocally and navigate through self-generated gradients remains largely unexplored. Here we introduce a theoretical framework for the self-organized chemotaxis of heterogeneous cell populations. We find that relative chemotactic sensitivity of cell populations controls their long-time coupling and co-migration dynamics, with boundary conditions such as external cell- and attractant reservoirs substantially influencing the migration patterns. We further predict an optimal parameter regime that leads to robust and colocalized migration. We test our theoretical predictions with in vitro experiments demonstrating the co-migration of different immune cell populations, and quantitatively reproduce observed migration patterns under wild-type and perturbed conditions. Interestingly, immune cell co-migration appears close to the predicted optimal regime. Finally, we incorporate mechanical interactions into our framework, revealing a phase diagram that illustrates non-trivial interplays between chemotactic and mechanical non-reciprocity in driving collective migration. Together, our findings suggest that self-generated chemotaxis is a robust strategy for emergent multicellular navigation.

## INTRODUCTION

Directional cell and tissue movement controls many key processes in development and disease such as morphogenesis, immune response and cancer invasion. While it is commonly assumed that this collective motion is steered by pre-patterned chemical or mechanical cues, only limited evidence has been gathered to support such long-range guidance in vivo [1]. In contrast, recent findings are unveiling the prominence of local, self-generated cues across a wide array of in vitro and in vivo settings [2–8], prompting a shift towards exploring controlled vs. *self-organized guidance* as alternative mechanisms of collective cellular dynamics [9].

Although recent experimental and computational studies have demonstrated self-generated chemotactic gradients as a robust guidance principle over large length scales [10], the primary focus has largely remained on the migratory patterns of homogeneous cell populations, or aimed at understanding phenotypic heterogeneity in bacterial migration [11–14]. However, directional cell migration frequently occurs through the coordination among different cell types, such as during processes like cell sorting [15], morphogenesis [16–20], wound healing [21], cancer invasion and growth [22, 23], and immune cell migration [8]. This ubiquity in *multi-component* cell communication raises fundamental questions about the role of self-organized guidance cues in navigating mixed cell populations towards specific targets.

From a theoretical standpoint, recent research has focused on the dynamics of chemotactic invasion with cell growth [24–26], and on clarifying the existence or stability of travelling concentration fronts of single populations [12, 25–29]. Some recent studies have also started to explore variability in cells’ chemotactic responses [11, 12, 14]. However, for a comprehensive understanding of multi-component navigation strategies via self-generated chemotaxis, it is essential to consider mixed cell types with distinct roles, such as chemotactic cell populations acting as sinks or sources in patterning [30].

Here, we propose a theoretical framework to explore the self-generated chemotaxis of mixed cell populations, concentrating on a *sink-sensor* system where two cell types are (asymmetrically) coupled through a diffusible chemoattractant. We find a rich spectrum of qualitatively distinct co-migration patterns, which can be understood as a function of few dimensionless parameters, such as relative chemotactic strength of different populations. In particular, our phase diagram predicts that robust and efficient co-migration (defined as both cell populations migrating at the same speed and being spatially colocalized) occurs in a specific region of parameter space, characterized by their relative chemotactic strength being close to unity. We also explore the role of interactions with external cell and attractant “reservoirs” in the system, taking two limits of initially fixed and renewing attractant levels, and find that these give rise to qualitatively different collective dynamics. Interestingly, we find that cell populations migrate as travelling waves only in the absence of an external attractant reservoir, where self-generated gradients remain sharp enough to facilitate robust front propagation. We furthermore the-oretically predict the propagation velocity and density profiles of coupled cell populations, and quantitatively test these predictions using different controlled in vitro migration assays with two mixed immune cell populations.

## RESULTS

### Continuum model of chemotactic cell mixtures

To explore the migration dynamics of heterogeneous cell populations, we turn to a continuum modeling approach adequate to describe cell and chemoattractant dynamics in a coarse-grained framework, extending the seminal work of Keller & Segel [31]. We focus on a minimal model for two distinct populations of chemotactic cells types: (i) *Consumer-sensor* cells that can actively modulate as well as sense gradients of the chemoattractant, which we will call *consumer* cells in short, and (ii) *sensor* cells that only sense and respond to it, with densities denoted by *ρ*_*c*_ and *ρ*_*s*_, respectively (see Fig.1A for an illustration). Effectively, this means that the first cell population is able to perform self-generated chemotaxis on its own, while the second population needs to “surf” on the gradient generated by the first one. The density evolutions of the cells are determined by an advection-diffusion equation (see Supplementary Information (SI) for more details):

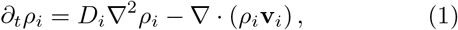

with the chemotactic drift velocity

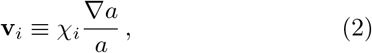

where *D*_*i*_ and *χ*_*i*_ denote the diffusion and the chemotactic coefficients of the cell type *i* = *s, c*, and *a* represents the chemoattractant concentration. Note that, this choice of logarithmic response represents the simplest form of a Weber-Fechner type chemotactic sensing, in the absence of more detailed information on the receptor kinetics of cells [14, 29, 32] (see SI for a brief discussion on alternative response functions).

The time evolution of the attractant *a* is determined by its diffusion and internalization by the consumer cell population. In the absence of an external reservoir, attractant density then follows:

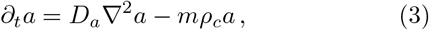

where *D*_*a*_ represents the diffusion coefficient of the chemoattractant, and *m* is the uptake rate of chemoattractant molecules by the consumer cells. The consumer cell population actively shapes the attractant profile, thereby controlling the chemotactic migration of the sensor cells. This effective interaction through an attractant field provides an asymmetric coupling between the two cell populations, reminiscent of chemically active matter mixtures with non-reciprocal interactions [33].

**FIG. 1.**
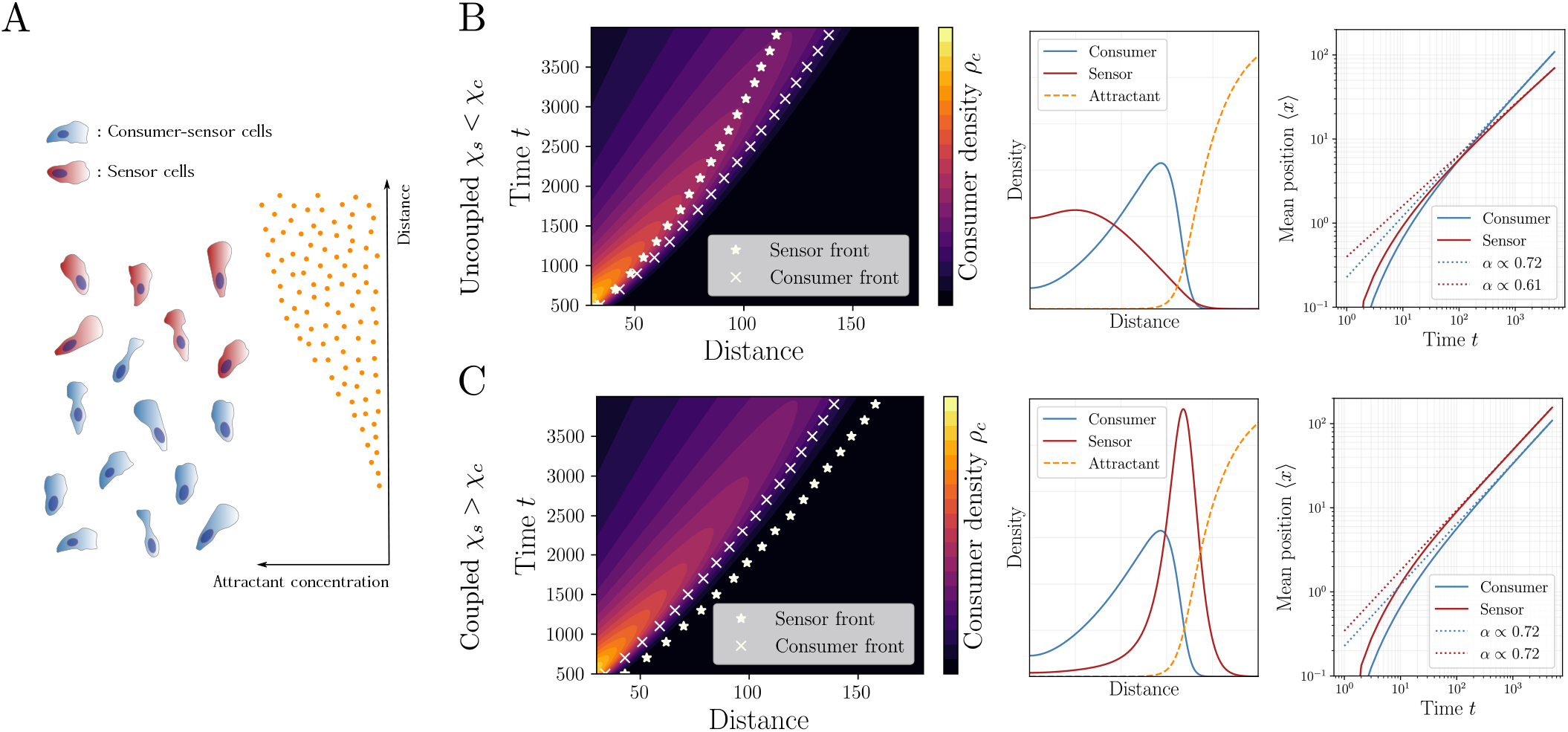
Coupled chemotaxis of mixed cell populations interacting through a locally modified attractant concentration. (A) Schematics of the theoretical model. Chemoattractant concentration (illustrated on the right) is dynamically shaped by *consumer-sensor cells* (blue) that can locally internalize the ligand, and preferentially migrate towards higher chemoattractant concentration. *Sensor cells* (red) only read off and respond to the local attractant gradient, but cannot modulate it. (B) If the rescaled chemotactic coefficient 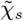 of the sensor cells is smaller than the coefficient 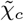 of the consumers, the sensor cells cannot follow the migratory front of the consumer cells, leading to the breakdown of their effective coupling through the attractant. (Left) Kymographs show that the sensor-cell front (star symbols) falls behind the consumer-cell front (crosses) for large times. (Middle) Density profile of the sensor cells (red) does not exhibit a well-defined peak and is confined to the back of the consumer cells (blue). (Right) The long-time behavior of the mean position as given by the relation ⟨*x*⟩ ∝ *t*^*α*^ shows that the asymptotic exponent of sensor cells attains a smaller value than that of the consumer cells, i.e. *α*_*s*_ < *α*_*c*_. (C) If the rescaled chemotactic coefficient of the sensor cells is larger than that of the consumers, i.e. for 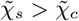, sensor cells migrate robustly ahead of the consumer cell front over long distances. (Left) Kymographs show that the sensor front is coupled to the consumer-cell front over large times. (Middle) Spatial profiles of cell populations exhibit coupled propagation of well-defined density peaks. (Right) Long-time scaling of the mean positions shows that the scaling exponents *α* of consumer and sensor cells are superdiffusive (i.e. *α* > 0.5) attaining similar values *α*_*s*_ ≃ *α*_*c*_ ≃ 0.7, which indicates their asymptotic coupling.

After rescaling time and space by *t* →*ηt*′, *x* → 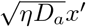 in Eqs.(1-3), where 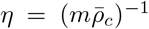 with a reference consumer density 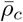, and dropping the primes for clarity, we obtain the nondimensional system of equations

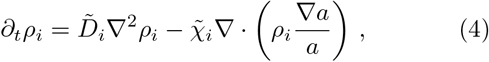

and

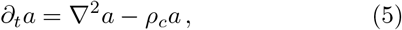

where 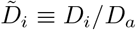 and 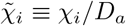 are the rescaled diffusion and chemotactic coefficients. We can now numerically evaluate Eqs.(4-5) for an initially uniform chemoattractant distribution over space, and with spatially localized initial cell density profiles to focus on their patterning dynamics driven by self-generated gradients, see SI for details on the numerical implementation.

### Co-migration of cell populations is regulated by their relative chemotactic response

An immediate question arises when we consider the coupled migration dynamics of the chemotactic cell populations: Can the self-generated guidance field provide a robust mechanism for sustained, long-range co-migration beyond diffusive or random motility? For a single cell population of consumer cells, a sufficiently large chemotactic coefficient 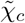 should allow for such a migration pattern, as observed in experiments [2, 3, 5, 8]. In agreement with these observations, we found that in the *chemotactic regime* with 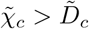, consumer cells propagated over long times with a well-defined density peak (see Supp.Fig.S1A). Longtime behavior of the mean position as determined by 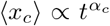 typically had an exponent *α*_*c*_ > 0.5, indicating super-diffusive motion (see Supp.Fig.S1B). Including the sensor cells in the system, we can now control their motility by tuning two parameters: Their (i) rescaled diffusion constant 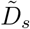, or (ii) rescaled chemotactic coefficient 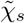. We first tested variations in 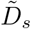 and found that this only led to uncoupled “spreading” of the sensor cell density without a large influence on the exact shape and position of the front (see Supp.Fig.S1C). In the following, we therefore decided to fix 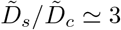 to consider generically “fast sensor” populations (as this also corresponded to the experimental data on immune cells as discussed below and in SI) and explored the remaining parameter space. Varying the relative strength of chemotaxis revealed the requirements for robust co-migration: When the chemotactic coefficient of the sensor cells is smaller than that of the consumers, i.e. 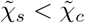, the sensor cells cannot keep up with the migratory front of the consumer cells and “fall behind”, with a mean position ⟨*x*_*s*_⟩ < ⟨*x*_*c*_⟩ and vanishing density peaks, see Fig.1B. In the chemotactic regime of consumer cells (i.e. for 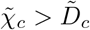), we found a tightly coupled propagation pattern only when 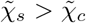, i.e. when the sensor cells have a stronger chemotactic sensitivity than the consumer cells. In this case, sensor cells have the potential to be faster than consumers, but cannot move arbitrarily far ahead of the consumer peak, as they then reach regions of flat chemoattractant gradients. Thus, in this parameter region, the sensor cell population exhibited a sustained density peak localized ahead of the consumer cells, see Fig.1C. For 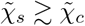, the coupled co-migration of the two cell populations was also reflected in their scaling exponents *α*_*c*_ ≃ *α*_*s*_. This is in contrast with the uncoupled regime 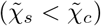, where the scaling exponents generically fulfil *α*_*s*_ < *α*_*c*_. Importantly, we also checked that these features were conserved for the case 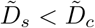 (see Supp.Fig.S1D).

### Consumer-sensor separation is controlled by chemotactic sensitivity

Next, we wanted to understand more quantitatively how the separation between the two cell populations is regulated as a function of control parameters. Can for instance sensor cells that exhibit fast random motility spread arbitrarily ahead of the consumer population? To explore this, we first systematically varied the chemotactic ability of the consumer cells 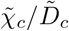 and the relative chemotactic sensitivity 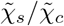 of the two populations. Examining the long-time ratio of the mean position of cell densities 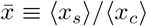 displays several distinct regimes for the co-migration dynamics, see Fig.2. (i) For sufficiently chemotactic consumer cells with 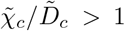, weak sensors 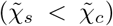 lead to uncoupled migration as discussed above, where the mean position ratio fulfils 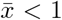 (see Region 1 in Fig.2A). Increasing the relative chemotactic strength 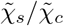 leads to larger ratios, with a crossover of 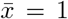 at 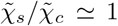. However, in the coupled regime, the position ratio saturates for arbitrarily large 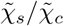, highlighting that the sensor cell propagation is controlled by the consumers (see Region 2 in Fig.2A). Interestingly, increasing the diffusion coefficient ratio 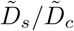, i.e. the random motility of the sensor cells, is only beneficial for the sensor cell population in the uncoupled regime with 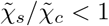, while for 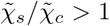 it has a negligible effect (see Fig.2B). (ii) For weakly chemotactic consumer cells with 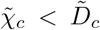, the phase diagram displays two additional regimes: In the region 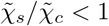, the mean position ratio exhibits 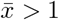 while not exceeding the upper bound 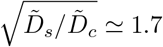 dictated by pure diffusion (see Region 3 in Fig.2A). Indeed, for identical sensor and consumer diffusion coefficients, position ratio in this region remained within 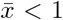 (see Supp.Fig.S2A). For larger relative chemotactic sensitivities, i.e. for 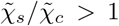, the position ratio can attain values larger than 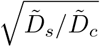, unlike the saturation behavior we found in the chemotactic regime 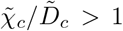 (see Region 4 in Fig.2A).

**FIG. 2.**
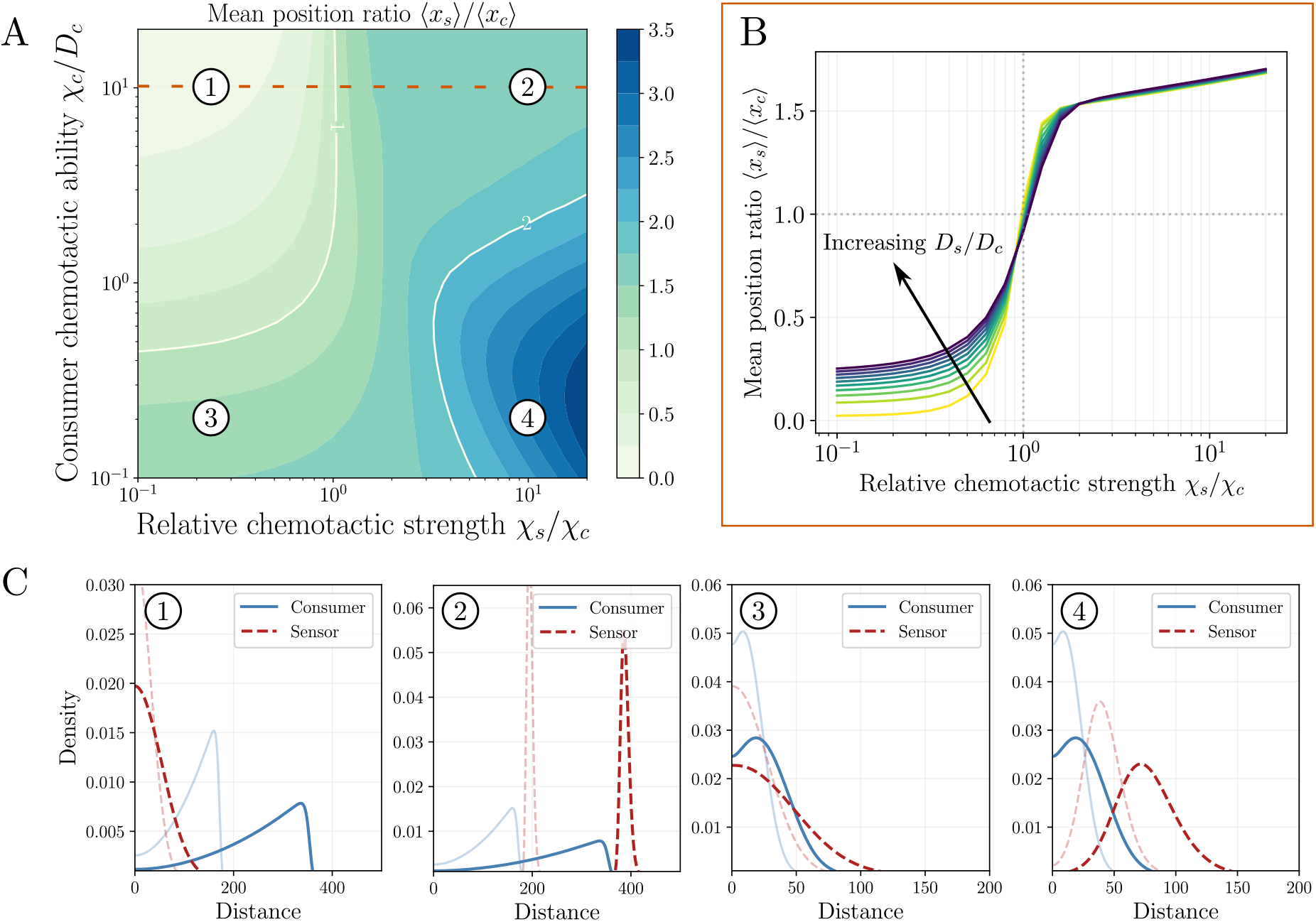
Relative positions and migration patterns of cell populations are controlled by their chemotactic sensitivity. (A) Long-time mean position ratio 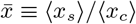 (color-coded) as a function of the rescaled chemotactic coefficient ratios 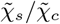 and relative chemotactic prowess of consumer cells, as controlled by the ratio 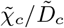. Dashed line (orange) corresponds to 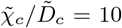 as plotted in (B). White contour lines outline the parameter regimes with 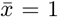 and 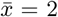. For consumer cells with a small 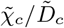 ratio, the sensor cell position can exceed the bound 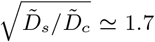 dictated by diffusion as 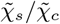 increases. (B) For sufficiently chemotactic consumer cells, e.g. with a chemotaxis-to-diffusion ratio 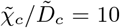, the mean position ratio 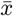 of consumer and sensor cell population densities exhibits a bounded increase as a function of chemotactic coefficient ratio 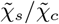, indicating that the sensor-cell position is controlled by consumer cells. Increasing the rescaled diffusion coefficient 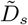 of the sensor cells (different colored lines) changes the ratio predominantly in the uncoupled regime for 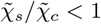. (C) Exemplary cell density profiles from different parameter regions plotted in (A) show the emergence of front- and pulse-like propagation patterns. Solid and dashed lines depict the profiles of consumer and sensor cells, respectively, at different time points (shaded lines).

A closer look at the density profiles in the four regions allowed for more insight into the propagation dynamics: For sufficiently chemotactic consumer cells, the chemotactic response of the sensor cells is a key parameter to tune their density profile and relative position with respect to the consumer cell population, with a sharp transition from uncoupled to coupled co-migration around 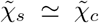. For large 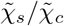 (e.g. Region 2) this coupling can then lead to the formation of a well-defined “sensor-cell pulse” ahead of the consumer cell front (see Fig.2C). When the consumer cell population is in the weakly chemotactic regime with 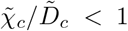, the sensor cell population can migrate transiently in a pulse-like manner at the diffusive tail of the consumers for large 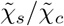 (e.g. Region 4).

Time evolution of the density patterns furthermore reveals that these are slowly decaying profiles (see shaded lines in Fig.2C), although their shapes are approximately scale-invariant (see Supp.Fig.S2B). In fact, the minimal system described by Eqs.(4-5) cannot exhibit travelling wave profiles in a strict sense [34] and the solutions will generically be “wave-like” profiles with slowly decaying amplitudes and velocities [35]. The coupling behavior shown in Fig.2 nevertheless remains preserved for sufficiently large times because of the slow decay dynamics (see Supp.Fig.S2C for a comparison with different time points.). The absence of travelling waves in a strict sense can be intuitively understood as follows: In the back of the migrating front (i.e. for small *x*) the uptake term − *ρ*_*c*_*a* is dominated by a small attractant concentration *a*, which leads to a shallow gradient. For conserved cell density, the chemotactic drift *χ* ∇ log(a) then cannot balance the diffusive leakage of cells in the back [36]. Therefore, boundary conditions such as cell conservation in the system and attractant kinetics should have a direct influence on the migration dynamics, as they will directly modulate the cellular responses behind the front.

### Influence of cell influx on the migration dynamics

To explore the effect of external interactions on the migration dynamics in greater detail, we turned to a simple modification where we allowed cell influx through the boundary at *x* = 0, with constant flux ∂_*x*_*ρ*|_*x*=0_ = *γ*, see Fig.3A for an illustration. Strikingly, a small nonzero rate *γ* led to the formation of travelling waves with a constant velocity: The consumer cell density showed an initial density peak, which decayed into a wave front profile with a constant density at the back, see Fig.3C, left panel. In the coupled regime with 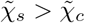, the sensor cell profiles attained a density peak ahead of the consumer cells, with a conserved separation between the two cell fronts (see Fig.3D, left panel). Kymographs of cell densities showed that the velocities of the cell fronts (as quantified by the half-maximum of peak densities) were strictly matched with 𝒱_*s*_ = 𝒱_*c*_ for 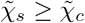, see Fig.3E, top panel. When the sensor cell chemotactic coefficient was smaller than that of the consumer cells, however, sensor cells again “fell behind” the consumer cell population with a decaying speed and density profile (see Supp.Fig.S3A-B). Interestingly, we found travelling waves for *both* cell populations even when the boundary influx *γ* was only nonzero for the consumer cell population, suggesting that consumer influx is sufficient for the efficient co-transport of sensor cells with a constant speed.

**FIG. 3.**
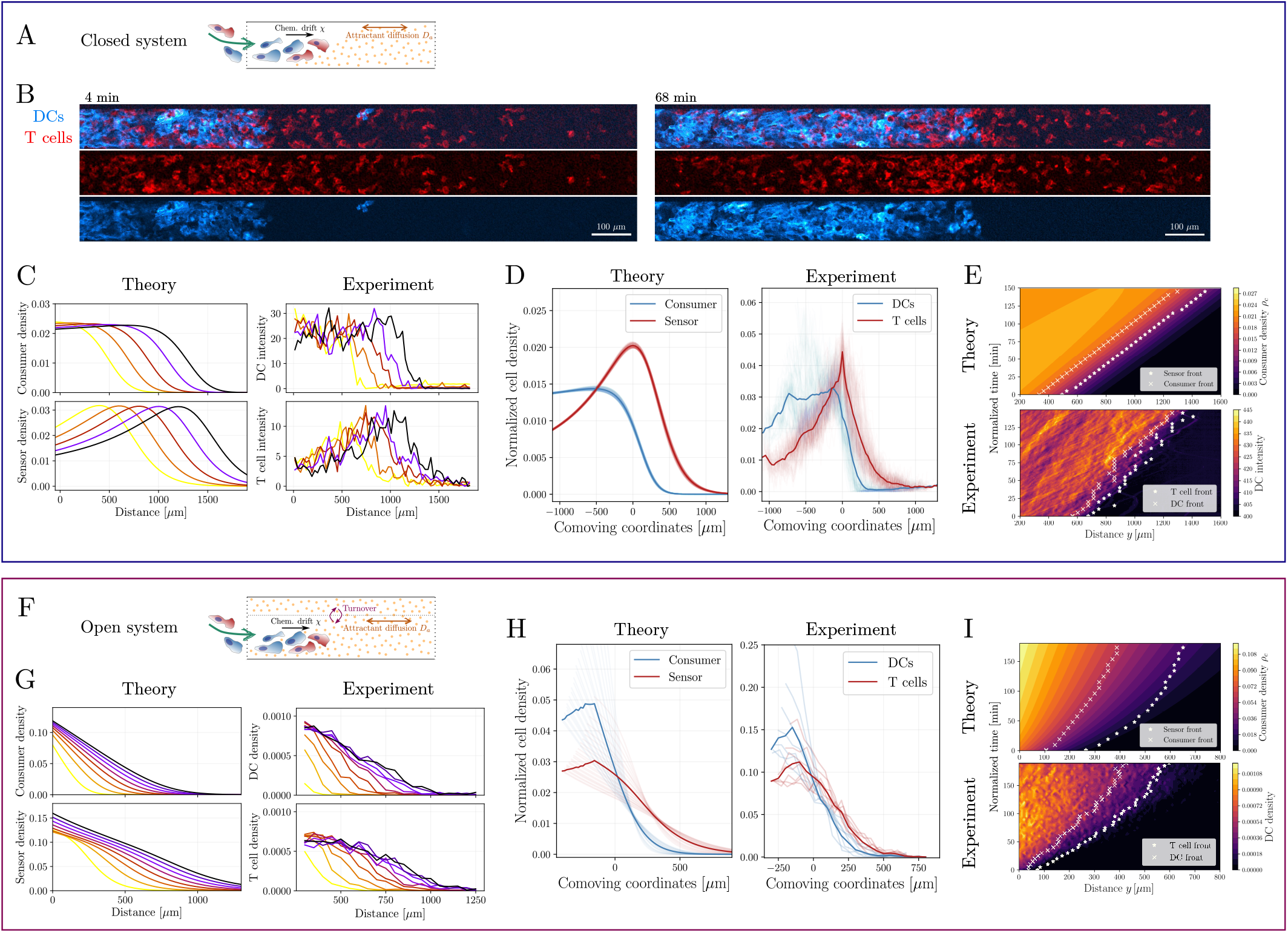
Comparison with in vitro experiments on collective migration of dendritic cells (DCs) and T cells. (A) Illustration of a *closed system*, where chemoattractant kinetics are described by its diffusion (with coefficient *D*_*a*_) and interaction with the migrating cells that locally consume the attractant and experience a chemotactic drift. Cells enter the channel from the left boundary. (B) Microscopy images from the microfluidic channel experiments with labelled DCs (blue) and co-migrating T cells (red) at *t* = 4min (left) and *t* = 68min (right). Scale bar indicates 100*µ*m. (C) Spatiotemporal cell density profiles predicted by the analytical model (left) agree well with the coupled propagation of front- and pulse-like densities as inferred from DC- and T cell intensities in experiments, respectively (right). Color code indicates different time points. (D) Normalized densities at different time points (shaded lines) overlaid in the reference frame comoving with the sensor-(left) or T-cell (right) peak. Density profiles are qualitatively conserved over time with a well-defined separation between the fronts. Kymographs showing the theoretical predictions on consumer migration (top) and experimental data on DC migration (bottom) over time. Cell density fronts are represented for both the consumer/DC (crosses) and the sensor/T cell (stars) population. Front positions over time indicate a constant speed of 𝒱 ≃ 6 *µ*m*/*min with which both cell populations migrate. Illustration of an *open system*, where a chemoattractant reservoir (delineated by the dotted line) enables in- and outflux of attractant molecules, leading to an effective turnover kinetics. (G) Analytical model (left) predicts spatiotemporal profiles without clear density peaks for both populations (left), as observed in experiments (right). (H) Cell densities in the comoving frame of the mean sensor cell (T cell) position display a non-conserved profile over time (shaded lines). (I) Kymographs showing the theoretical predictions on consumer migration (top) and experimental data on DC migration (bottom) over time. Density fronts are represented for both the consumer/DC (crosses) and the sensor/T cell (stars) population. Front positions over time indicate a slowly decaying speed both for theory and experiment. Experimental data plotted in (G-I) from [8].

### Comparison to experimental data on the self-organized co-migration of immune cell populations

To test our theoretical prediction on the coupled migration patterns, we turned to a minimal experimental setup with heterogeneous immune cell populations. We focused on the collective migration of dendritic cells (DCs) and T cells, as we had recently observed qualitatively their co-migration patterns based on self-generated gradients of the chemoattractant CCL19 [8]. To closely match the coarse-grained description of the cell density and chemoattractant evolutions governed by Eqs.(4-5), we designed a microfluidic channel system with two holes to control both the cell and attractant influx. Because there is no external chemoattractant reservoir in this setup, it represents a “closed” system with regards to attractant kinetics. After an equilibration period to allow for a uniform distribution of CCL19 in the channel, we introduced mixtures of DCs and T cells to the microfluidic device through one end of the channel. We observed that both cell populations continuously entered the channel for time periods exceeding 3-4 hours, see Fig.3B. Strikingly, we found that DC population migrated as a front with an approximately constant density profile in the back, while T cells showed a clear density peak ahead of the DC front, as we predicted theoretically, see Fig.3C. Overlaying the density profiles at different times in a comoving coordinate system centered at the T cell peak, we found that both density profiles were well-maintained over time, with the two fronts separated with a characteristic length scale of approx. 150-200 *µ*m, see Fig.3D. Furthermore, the kymographs obtained from the experiments confirmed the theoretical prediction on velocity matching between the two populations (see Fig.3E). To test our prediction on the breakdown of velocity matching in the uncoupled regime (i.e. for the case 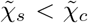), we then looked at the co-migration of DCs with CCR7-knockout (KO) T cells (which do not have the capacity to sense the ligand CCL19) in microfluidic channel experiments. We found that CCR7-KO T cells (i) did not show any density peaks, (ii) were not able to migrate ahead of the DC population, and (iii) their speed was notably smaller than that of the DC front (see Supp.Fig.S5A-D). Interestingly, even for WT T cells we found that their propagation speed was notably reduced when they migrated alone (in the absence of the DCs) either in a gradient or uniform field of CCL19 (see Supp.Fig.5E).

### Prediction of the front velocity and parameter estimates

We then sought to constrain model parameters via two independent experimental setups. First, using trajectory datasets from under-agarose assays with only DCs migrating without a chemoattractant [8], we estimated their diffusion coefficients to be around *D*_*c*_ ≃ 350 *µ*m^2^*/*min, leading to a rescaled diffusion coefficient of around 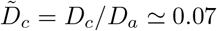 with a chemoattractant diffusion coefficient *D*_*a*_ ≃ 86 *µ*m^2^*/*s (as we measured for CCL19) [8], see Supp.Fig.4A-C. Second, we used trajectories from the under-agarose assay for mixed populations of DCs and T cells with CCL19, and calculated the variance of velocity distributions for both populations, as the velocity fluctuations are proportional to their random motility coefficient. The comparison of this variance allowed us to approximate the ratio of their diffusion coefficients to be around *D*_*s*_*/D*_*c*_ ≃ 3.5 (see Supp.Fig.4D and SI Section S3 for details on the diffusion coefficient estimation).

Interestingly, the observation of a traveling cell front allowed us to theoretically predict the spatial shapes of the density profiles, as well as the propagation velocity 𝒱 using simple arguments in the co-moving frame of co-ordinates *z* ≡*x* − *𝒱t*. Indeed, we predict that both the consumer and sensor cell populations should display exponentially decaying density profiles at the propagating edge with *ρ*_*i*_ ∝ exp(− *ζ*_*i*_*z*), with a decay length *ζ*^−1^ related to the front velocity by 𝒱 ∝ *D*_*i*_*ζ*_*i*_. This relation means that the ratio of decay lengths is simply proportional to the ratio of diffusion coefficients between the consumer and sensor cell populations (*D*_*s*_*/D*_*c*_ = *ζ*_*c*_*/ζ*_*s*_, which we confirmed numerically, (see Supp.Fig.S3C). Importantly, we then used this relation to compare the experimental density profiles of DC (consumer) and T cell (sensor) populations in the comoving frame, and from their decay lengths obtained a value of *D*_*s*_*/D*_*c*_ ≃ 2.5 (see Supp.Fig.S4E), closely aligning with independent estimates described above.

Furthermore, we numerically observed that the consumer cells exhibit a spatially flat profile behind the front with a constant density 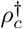, with the chemoattractant concentration showing an exponential decay approximated by *a* ∝ exp(*λz*) in the back of the front. Using this information, we integrated the system of equations in the comoving frame and found that the decay length scale of the chemoattractant and the front velocity are linked by the relation 𝒱 = *χ*_*c*_*λ*. In dimensional units, we could then obtain a simple expression for the front velocity (see SI Section S1 for the derivation):

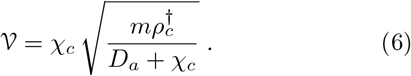

This relation suggests that the front velocity scales as 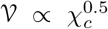, and depends on an effective time scale of chemoattractant internalization 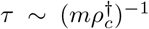 fixed by the rate *m* and bulk consumer density 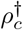, while similar expressions have been found under different assumptions [25, 28]. We then tested this prediction for the front velocity numerically by varying the chemotactic co-efficient *χ*_*c*_, and found good agreement with Eq.(6) (see Supp.Fig.S3D).

These analytical results provided us a robust method to fit the rescaled chemotactic coefficients 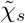 and 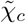 in our system. Qualitatively, we noted that for sufficiently chemotactic consumers 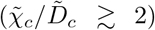, where a well-defined traveling front can be formed, the relative chemotactic strength should not be larger than 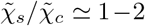 to prevent a sensor cell accumulation far ahead of the consumers. Additionally, for 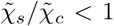, we transition into the uncoupled regime where the sensor cell population cannot keep up with the consumer front, which together strongly constrained the range of possible chemotactic strength ratios.

We then used a sufficiently large chemotaxis-to-diffusion ratio of 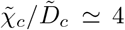 for consumer cells to obtain a well-defined wave front, and fitted the timescale 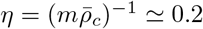 min which provided a good quantitative agreement between theory and experiment for several distinct features (as displayed in Fig.3): (i) the scaling of the density profiles at both front and back agreed closely with the experimental data, (ii) theoretical values obtained for the separation between the two populations were about 200 *µ*m vs. about 150 *µ*m in experiments, and (iii) the theoretical prediction for the front velocity with a value of 𝒱 ≃ 6 *µ*m*/*min quantitatively matched the experimental front speeds. Finally, we used the aforementioned values for *χ*_*c*_, *D*_*a*_, and only by fitting the time scale 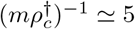 min in Eq.(6), we obtained an analytical estimate of 𝒱 ≃ 6.7 *µ*m*/*min, in close agreement with both experiment and the numerical results.

### Influence of chemoattractant turnover on the migration dynamics

Next, we asked whether modifying the chemoattractant kinetics might influence the migration patterns in addition to the boundary influx of cells. Indeed, Eq.(5) suggests that a turnover of attractant density, e.g. arising from an external reservoir, should directly influence the dynamics of the gradient. We thus implemented a minimal model for the attractant kinetics to describe additional decay and influx terms, see Fig.3F for an illustration and SI for details. We found that numerically solving this “open system” led to strikingly different migration patterns: First, density profiles exhibited monotonically decaying shapes without well-defined peaks in a large region of the parameter space. In fact, sensor cell “pulses” only existed for relatively large chemotactic coefficients and consumer cells never exhibited pulse-like density peaks (see Supp.Fig.S6). Second, we could not find any parameter regime that led to the formation of travelling waves. Density profiles generically evolved with slowly decaying velocities. Third, even for very large chemotactic coefficients, the long-time front propagation dynamics showed diffusive scaling exponents around *α* ≃ 0.5, unlike the travelling waves of the closed system.

To test these theoretical signatures, we turned to experimental data on the collective migration of DCs and T cells in the under-agarose assay, as we had qualitatively explored before [8]. In this system, a large reservoir of the chemoattractant CCL19 is in constant contact with the confined quasi-2D space where the cells migrate (see SI for further discussion). Plotting the cell density profiles over time, we found that the densities exhibited monotonically decaying profiles as predicted by the theory, see Fig.3G for a comparison. Overlaying the profiles at different times, we also found that T cells migrated ahead of the DC population at all times with a length scale of the same order of magnitude as in the microfluidic channel, see Figs.3H, but with an increasing separation of cell fronts over time, see Fig.3I. This behavior matches with the theoretical prediction using a chemotactic strength ratio of 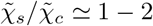 that we had previously estimated for the closed system, although their absolute values in the open system are larger. Finally, the long-time propagation dynamics of density fronts showed that both populations migrated with decaying velocities over time, as represented in the kymographs in Fig.3I. Overall, the analysis of this minimal open system indicates that attractant kinetics can indeed lead to radically different migration dynamics for chemotactic cell populations.

### Relative chemotactic sensitivity controls colocalized migration of cell populations

Having explored the coupled migration regimes in different setups, we then wished to go back to the minimal system described by Eqs.(4-5) and ask whether we could quantitatively measure the degree of overlap or *colocalization* between distinct populations. In addition to co-migration, the question of colocalization of heterogeneous cell populations is important particularly to explore conditions for possible physical interactions between different immune cell types, as they need to communicate in close proximity during an immune response [37]. To quantify this, we first defined a colocalization index using the Jensen-Shannon divergence (JS) of spatial density profiles interpreted as probability densities (see SI Section S5 for details), which provides a simple metric for similarity or overlap of cell densities as *ϕ* ≡ 1 − *D*_*JS*_, where *D*_*JS*_ denotes the Jensen-Shannon divergence. With this definition, *ϕ* = 1 corresponds to complete colocalization while *ϕ* = 0 indicates zero overlap of densities. Interestingly, this simple metric already outlined two distinct regions in the phase space where consumer migration occurs (for sufficiently large 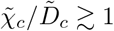), but with impaired colocalization, see Fig.4. Intuitively, in the uncoupled regime 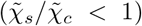, colocalization is low as the sensor cells in this case fall behind the propagating consumer front. However, in the *strongly coupled* regime 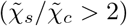, the sensor density peak is too far ahead of the consumers, which also results in minimal density overlap. Furthermore, while random cellular diffusion (controlled by 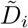) is detrimental for robust chemotactic front propagation, it is in fact beneficial for colocalization, as very small diffusion coefficients result in very sharp density peaks, and thus small density overlaps. This analysis thus predicted that each parameter of the system exhibited tradeoffs, while acting to optimize for both co-migration and colocalization. Notably, the values which fitted best our experimental data were located in the intermediate and “optimal” region in the coupled regime of the phase diagram where 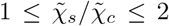 and 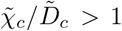. Finally, while we used identical diffusion coefficients for sensors and consumers in Fig.4, changing to experimentally inferred parameter estimates for the diffusion coefficients only modified the colocalization in the diffusively uncoupled regime, leaving the remaining phase diagram unchanged (see Supp.Fig.S7).

**FIG. 4.**
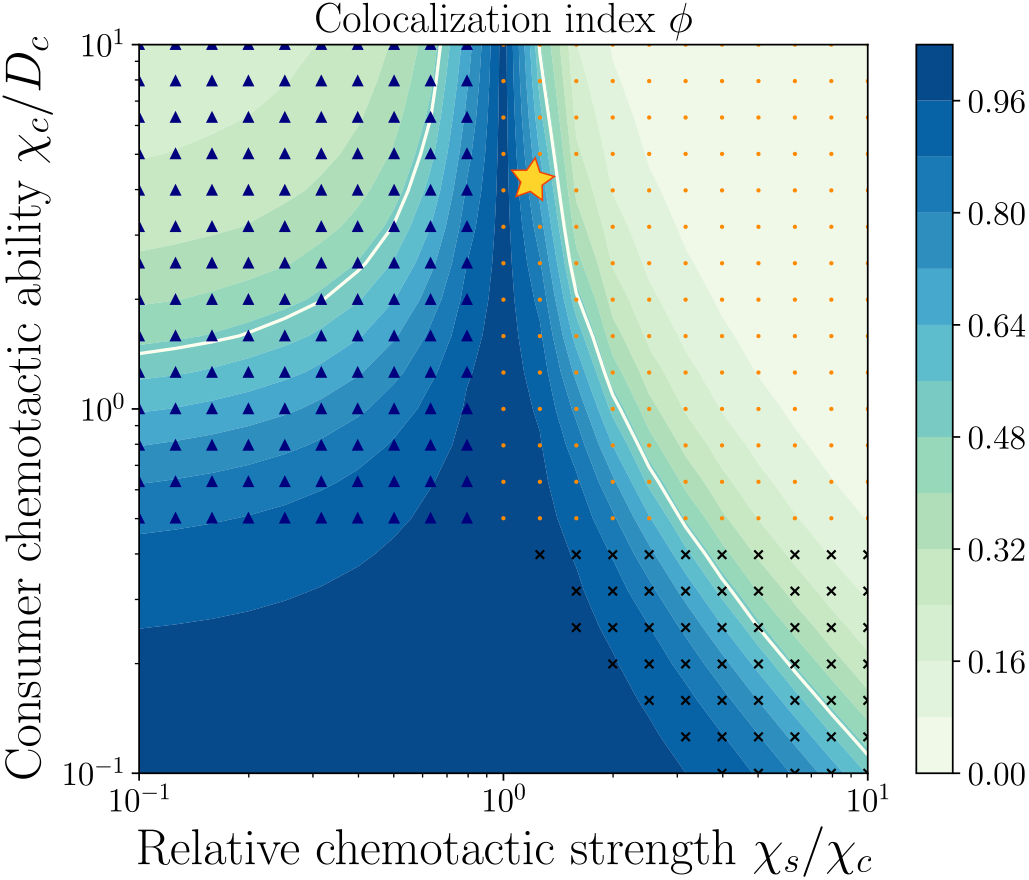
Phase diagram of colocalization of different cell populations indicates an optimal regime for coupled migration. (A) Colocalization index (color-coded) defined as *ϕ* ≡1 − *D*_*JS*_, using the Jensen-Shannon divergence (*D*_*JS*_) of density distributions for sensor and consumer cells. Contour lines with *>* 0.5 indicate sensor cells being transported by consumers with enhanced colocalization. For sufficiently chemotactic consumer cells, i.e. for 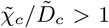, the colocalization index displays a non-monotonic behavior with increasing 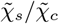, with an optimal region for 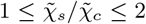. Parameter values used for the comparison with the experiments are 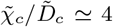 and 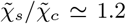 (star symbol). Distinct regions of density patterns are highlighted with markers for (i) only pulse-like consumer cells (triangles), (ii) pulse-like consumer and sensor cells (dots), (iii) no peaks (no markers), and (iv) only pulse-like sensor cells (crosses). Rescaled diffusion coefficients used here are 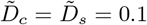.

### Influence of non-reciprocal mechanical interactions

We next asked whether migration patterns driven by self-generated chemotaxis are affected by mechanical interactions between heterogeneous populations, which in all generality can be non-reciprocal, e.g. if population A repels population B while population B attracts population A [38]. To address this, we considered a minimal model incorporating an advective flux term into the density evolution equations, where each population responds linearly to the density gradient of the other:

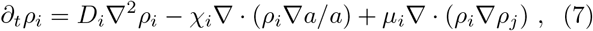

with *i* ≠ *j*, and *µ*_*i*_ represents the mechanical advection parameter for the *i*-th cell population (*i* = *c* for consumers, *i* = *s* for sensors). Positive values (*µ >* 0) indicate repulsion (cells being “pushed”), while negative values (*µ <* 0) represent attraction (cells being “pulled”) - see SI Theory.

Numerical simulations revealed several non-trivial features that emerged from the interplay of mechanical and chemotactic coupling. First, we found that the mean position ratio, 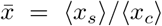, showed significant sensitivity to mechanical coupling for weakly chemotactic consumers (i.e. for small *χ*_*c*_*/D*_*c*_). In this regime, increasing sensor advection (large *µ*_*s*_) while reducing consumer advection (small *µ*_*c*_) led to a marked increase in 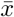 (Fig. 5A). Thus, this effect emerges specifically from non-reciprocal nature of the system, as quantified by the difference between both mechanical couplings Δ*µ* ≡*µ*_*s*_ − *µ*_*c*_. This became less pronounced as consumer chemotaxis strengthened (*χ*_*c*_*/D*_*c*_ *>* 1), where 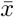 converged to predictions for mechanically non-interacting populations (Fig.5B, Supp.Fig.8).

**FIG. 5.**
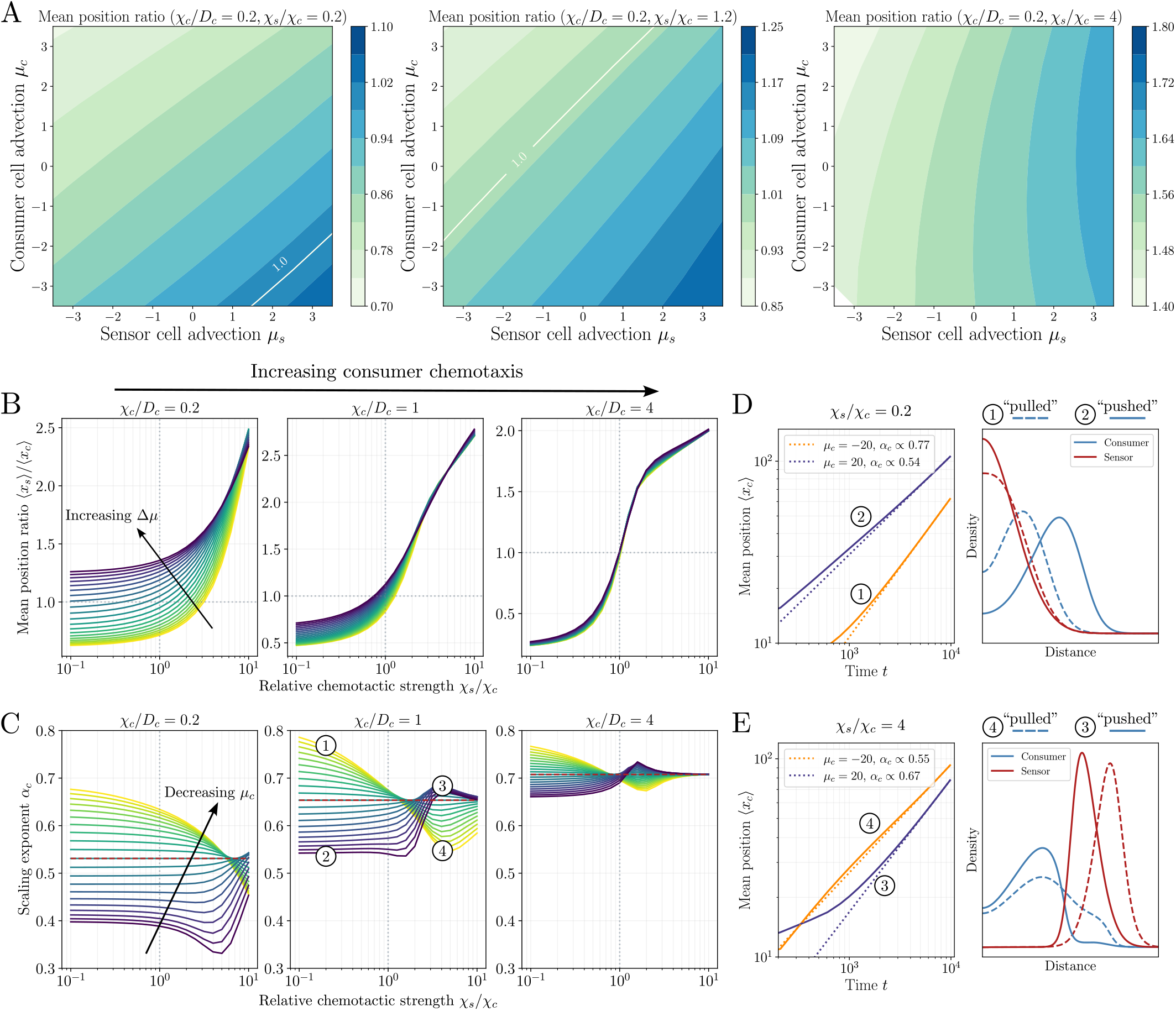
Influence of non-reciprocal mechanical interactions. (A) Mechanical coupling, controlled by parameters *µ*_*c*_ and *µ*_*s*_, influences the mean position ratio of cell populations in the weakly chemotactic regime of consumers (*χ*_*c*_*/D*_*c*_ ≃ 0.2). Variations in the coupling difference Δ*µ* = *µ*_*s*_ − *µ*_*c*_ significantly affect the mean position ratio for different sensor chemotactic strengths (increasing from left to right panels). (B) For weakly chemotactic consumers (leftmost panel), large positive Δ*µ* values (“pushed” sensors, “pulled” consumers) increase the relative positions. As the chemotactic ability of consumer cells increases (middle and right panel), the effect of mechanical coupling diminishes. (C) Scaling exponent *α*_*c*_ of the consumer cell population inferred from the long-time evolution of the mean position ⟨*x*_*c*_⟩ ∝ *t*^*αc*^ depends on the mechanical coupling parameter *µ*_*c*_, with most variability observed for small *χ*_*c*_ and *χ*_*s*_ (leftmost panel). Dashed line (red) indicates the predicted values for *µ*_*c*_ = 0. Intermediate consumer chemotaxis with *χ*_*c*_*/D*_*c*_ = 1 (middle panel) shows distinct regimes: weak sensor chemotaxis (*χ*_*s*_*/χ*_*c*_ < 1) increases *α*_*c*_ with consumer attraction (*µ*_*c*_ < 0, region 1) and decreases it with repulsion (*µ*_*c*_ *>* 0, region 2). For strong sensor chemotaxis (*χ*_*s*_*/χ*_*c*_ *>* 1), the trend reverses: repulsion increases *α*_*c*_ (region 3), while attraction decreases it (region 4). For strongly chemotactic consumers (*χ*_*c*_*/D*_*c*_ *>* 1, rightmost panel), mechanical coupling has a smaller effect on *α*_*c*_. (D) Evolution of mean positions (left panel) and density profiles (right panel) for weakly chemotactic sensors (*χ*_*s*_*/χ*_*c*_ = 0.2). Consumers with *µ*_*c*_ *>* 0 (“pushed”) exhibit more advanced mean positions than with *µ*_*c*_ < 0 (“pulled”). Density profiles show stronger overlap between sensors and consumers for *µ*_*c*_ < 0. (E) For strongly chemotactic sensors, consumer mean positions are more advanced for *µ*_*c*_ < 0 than for *µ*_*c*_ *>* 0 (left panel). However, for *µ*_*c*_ < 0, consumer density profiles lose their sharp fronts (right panel).

Second, to test whether mechanical coupling could mimic chemotactic dynamics for diffusive or weakly chemotactic consumers, we analyzed the long-time behavior of the consumer mean position 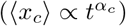 quantified by the scaling exponent *α*_*c*_. We found that *α*_*c*_ was primarily influenced by *µ*_*c*_, with the largest effects observed for small *χ*_*c*_ and *χ*_*s*_ (see Fig. 5C). For intermediate consumer chemotaxis (*χ*_*c*_*/D*_*c*_ = 1), distinct migration regimes emerged depending on sensor chemotactic strength: (i) When sensor cells are weakly chemotactic (*χ*_*s*_*/χ*_*c*_ < 1) and lag behind the consumer population, consumer attraction to sensors (*µ*_*c*_ < 0) increased the scaling exponent *α*_*c*_, while repulsion (*µ*_*c*_ *>* 0) decreased it. (ii) In contrast, for strongly chemotactic sensor cells (*χ*_*s*_*/χ*_*c*_ *>* 1) that migrate ahead of consumers, consumer repulsion by sensors (*µ*_*c*_ *>* 0) increased *α*_*c*_, while attraction (*µ*_*c*_ < 0) decreased it.

These trends reflect the system’s inherent non-reciprocity: For *µ*_*c*_ *>* 0 and weak sensor chemotaxis, consumers are “pushed from behind,” achieving greater mean positions while increasing their separation from sensor densities (Fig. 5D). This repulsion reduces the consumers’ ability to sense the attractant gradient efficiently. Conversely, for strong sensor chemotaxis, consumers are “pulled from ahead” (*µ*_*c*_ < 0), resulting in a broader density profile that locally flattens the attractant gradient and reduces the dynamic scaling exponent (Fig.5E). These findings thus highlight how mechanical interactions can modulate chemotactic co-migration patterns and sensing dynamics in regimes of weak or intermediate consumer chemotaxis.

## DISCUSSION

Self-generated chemoattractant gradients have recently emerged as an attractive mechanism for guiding cellular collectives over long distances in a robust and emergent manner [10]. In this work, we propose a theoretical framework demonstrating how this mechanism can also couple heterogeneous mixtures of cells without relying on direct intercellular interactions. We tested several key predictions of the model experimentally in the context of co-migrating dendritic and T cells. Our model quantitatively reproduces various features of their self-generated guidance, including the spatial shape of the density profiles, relative cellular speeds, and the influence of external reservoirs on the dynamics. Additionally, it identifies several qualitatively distinct modes of co-migration for different populations and highlights trade-offs in key model parameters. For example, for sensor cells to keep up with the propagating front of consumer cells, their chemotactic strength must be strictly larger. This typically results in a density peak of sensors ahead of consumers. However, if the chemotactic strength of sensors is too high, it leads to density peaks far from consumers, resulting in poorly colocalized migration. Similar trade-offs exist for cellular diffusion via random motility, which, while lowering overall chemotactic efficiency, increases the likelihood of cell colocalization. Interestingly, direct (non-reciprocal) mechanical interactions between cell populations primarily influence the migratory behavior in the limit of weak chemotaxis.

From a biological perspective, the physical interaction between DCs and T cells represents the central regulatory event in adaptive immunity. These encounters occur in the lymph node, where DCs enter via the lymphatic system and T cells via the blood vasculature [39]. It is firmly established that the commonly expressed chemokine receptor CCR7 ultimately guides both cell types into the same compartment, with a key role for the stromal cells in producing chemoattractants such as CCR19. However, the interaction between DCs and T cells is not a one-time encounter but rather a dynamic crosstalk where cells repeatedly meet and scan each other to integrate signals on the population level [40]. How this crosstalk is regulated remains poorly understood. Our findings propose a 10 simple mechanism via a self-generated chemoattractant field that facilitates sustained interactions and coupled migration of these distinct immune cell populations.

Collective long-range guidance of cell populations can emerge through various local interactions such as cell-cell repulsion, contact-inhibition of locomotion, polarity alignment [41], as well as mechanochemical coupling [42]. Although numerous modeling approaches including vertex and Voronoi models [43, 44], phase-field models [45], and continuum models e.g. based on non-local sensing mechanisms [46] have been applied to address the emergence of collective directionality, how heterogeneous cell populations with varying microscopic responses spatially pattern and control their co-migration remains to be an open question [30]. In particular, testing further the relative roles of chemical vs. mechanical guidance cues that regulate the interactions between heterogeneous cell mixtures will be a crucial step for decoding their patterning behaviour in different contexts.

Beyond chemoattractants, other types of self-generated gradients based on substrate rigidity or ECM remodelling have emerged in recent years [47–49]. These modes of guidance bear conceptual similarities with self-generated chemotaxis as cells modify their environment to encode a memory of their past trajectories, leading to their emergent directionality. While cell-driven changes to substrate mechanics or ECM composition are expected to produce more local cues than diffusible chemoattractant signals, we expect the core features of our proposed model to hold in these distinct scenarios. Here, interactions between sensors and consumers are intrinsically non-reciprocal due to their asymmetric coupling through the attractant, directly linking our model to recent advances in synthetic active matter on non-reciprocal systems. Incorporating mechanical interactions into our framework further suggests parallels with the collective behavior observed in chemically active mixtures of particles [33, 50] or travelling states of diffusive species where non-reciprocity enters via cross-diffusion [51].

Our theoretical model suggests a subtle mechanism to regulate the robust, long-range migration of heterogeneous cell populations in the absence of pre-patterned gradients. Such multicellular streams have been observed, for example, in CCR7-dependent trafficking of T cells in zebrafish [52] and their directional crawling between two compartments in the mouse spleen. In both scenarios, it seems unlikely that a fixed spatial gradient could guide the cells over distances of many millimeters, and rather suggests a role of distinct cell populations dynamically shaping the gradient, as demonstrated in the developing lateral line during zebrafish development [2, 53]. Our results on coupled migration patterns can provide insights for these observations and for further experimental studies such as in vitro motility assays in microfluidic devices [13], and to interpret large-scale live imaging of migrating heterogeneous cell populations [54].

## METHODS

### Cell culture

All cells used in this study were grown and maintained at +37C with 5% CO_2_ in a humidified incubator.

### Dendritic cell differentiation and maturation

DCs were generated from previously described LifeAct-GFP expressing conditionally HoxB8 immortalized hematopoietic progenitor cells ([55, 56]). LifeAct-GFP Hoxb8 cells were maintained in R10 medium (RPMI 1640 medium (Gibco, 21875-034) with 10% heat-inactivated fetal bovine serum (FBS) (Gibco, 10270-106), penicillin (100 U/ml)/streptomycin (100 *µ*g/ml) (Gibco, 15140-122), and 50 *µ*M *β*-mercaptoethanol (Gibco, 31350-010)) supplemented with 5% supernatant of an Flt3L-producing cell line and 1 *µ*M estrogen (Sigma-Aldrich, E2758). For DC differentiation, 3×10^5^ Hoxb8 precursor cells were cultured in 10 ml R10 supplemented with 20% of house-generated Granulocyte-macrophage colony stimulating factor (GM-CSF) hybridoma supernatant. On day 3 of differentiation, additional 10 ml R10 medium supplemented with 20% GM-SCF were added to the dish, on day 6 followed by replacement of half of the R10 medium with 20% GM-SCF. On day 9 DCs were harvested for maturation. DC maturation was induced by overnight incubation with lipopolysaccharide (LPS, 200 ng/ml) from Escherichia coli 0127:B8 (SigmaAldrich, L4516).

### T cells

T cells were isolated from spleens of male mTmG reporter [57] and CCR7 deficient mice [58] using the EasySep™ Mouse T Cell Isolation Kit (STEM-CELL Technologies, 19851) following manufacturers’ instructions and activated with anti-mouse CD28 (Invitrogen, 16-0281-85) and CD3e (Invitrogen, 16-0031-85) 1 *µ*g/ml coated to cell culture dishes. Activated T cells were cultured in R10 medium substituted with 10 ng/ml IL-2 (PeproTech, 212-12) for up to 14 days. Before migration assays CCR7 deficient T cells were stained with TAMRA (Invitrogen) 1:1000 in PBS for 10 min at room temperature, washed 2x with PBS right before the assay.

### Migration assays in microfabricated channels

Microfabricated polydimethylsiloxane (PDMS) devices were prepared as described previously [8]. Fabricated PDMS devices with straight channels (300 x 100 x 4,5 *µ*m) were flushed with R10 medium. For uniform chemokine assay, one of the reservoirs in the PDMS device was refilled with 1,25 ng/*µ*l of CCL19 (R&D Systems, 440-M3-025) supplemented R10 medium and allowed to equilibrate for 2-3 hours at 37 °C and 5% CO2. After the incubation, 2 *µ*l of pelleted DC and T cell mixture was added to the opposite side reservoir in the PDMS device. For T cell migration in chemokine gradient, 2 *µ*l of pelleted T cells were added at one side of the device and 1,25 ng/*µ*l CCL19 in R10 medium to the opposite side at the same time. The time-lapse imaging of migrating cells was performed by imaging every 30 s with an inverted Nikon Ti2E wide-field fluorescent microscope at 37°C with 5% CO2 for the duration of 4-6hours, using Nikon 10x objective (Plan Fluor 10x/0.3 DIC 1 N1 air PFS).

## Supporting information

Supplementary Theory Note

## AUTHOR CONTRIBUTIONS

Project conceptualization: M.C.U., E.H., and M.S.; Theoretical model: M.C.U. and E.H.; Experiments and data acquisition: Z.A.; Data analysis: M.C.U.; Writing, original draft: M.C.U. and E.H.; Writing, review and editing: All authors.

## ACKNOWLEDGEMENTS

We thank all members of the E.H. and M.S. groups for stimulating discussions. We thank the Imaging and Optics facility, the Pre-clinical and Lab Support facility of the Institute of Science and Technology for their excellent support and provided resources for the experimental research. In particular we thank Jack Merrin from the Nanofabrication facility who generated the microfabricated channel used in this study. This work received funding from the ERC under the European Union’s Horizon 2020 research and innovation program (grant agreement No. 851288 to E.H.).

