## Supplementary Theory Note for "Self-generated chemotaxis of mixed cell populations"

December 17, 2024

**4 Migration patterns in the open system** **15**

**5 Jensen-Shannon divergence for colocalization** **17**

**6 Mechanical interactions between cell populations** **20**

**7 Details on numerical solution of the PDEs** **21**

**8 List of Supplementary Movies** **22**

In this Supplementary Note, we provide additional details on the modelling approach, analytical predictions, parameter inference, numerical solution methods, as well as sensitivity analyses on how different model setups and parameter choices affect our results.

### 1 Details on the coarse-grained model

Continuum modeling approaches for chemotaxis can be generically formulated in the framework of a persistent and biased random walk, as it was introduced by Patlak [1]. Different microscopic mechanisms based on Fick's law, "space-jump" processes, or transport equations can be used to obtain coarse-grained continuum models for chemotaxis [2]. These different derivations lead to coupled PDE systems, as analyzed by Keller and Segel [3, 4], which can result in spatial patterning when the cells both produce and migrate up chemoattractant gradients. Although the original Keller-Segel system involved a single chemoattractant-modulating cell population together with the dynamics of the attractant, it is rather straightforward to introduce additional chemotactic cell populations in this framework. One can then ask whether the different cell types will be governed by a symmetric description in terms of (i) their chemotactic sensing/response, which controls their advective speed, and/or (ii) their interactions with the chemoattractant concentration, i.e. whether they act as producers or consumers of the attractant. In the context of bacterial chemotaxis, for instance, such *heterogeneity* in chemotaxis has been studied with respect to the chemotactic responses of different cell types [5, 6, 7, 8]. However, heterogeneity in terms of attractant modulation has remained largely unexplored. Here, we will focus on a Keller-Segel framework to describe mixed cell populations with distinct roles where we have *consumer-sensor* cells that both sense and shape, and *sensor* cells that can only sense attractant gradients. The simplest formulation of the coupling between these two cell populations via the chemoattractant is then defined by

$$\partial_t \rho_i = D_i \nabla^2 \rho_i - \nabla \cdot (\rho_i \mathbf{v}_i), \tag{S1}$$

with the drift velocity $\mathbf{v}_i \equiv \chi_i \nabla \log(a) = \chi_i \nabla a/a$, and the subscript $i = c, s$ describes the consumer or sensor population, respectively. The chemoattractant profile will be governed by diffusion with coefficient $D_a$ and its consumption by the consumer population with rate $m$:

$$\partial_t a = D_a \nabla^2 a - m \rho_c a, \tag{S2}$$

Note that the chemotactic drift velocity $\mathbf{v} = \chi \nabla \log(a)$ represents the simplest formulation of a Weber-Fechner type of attractant sensing (relative sensing), as discussed originally by Keller and Segel. Alternative formulations for the sensing are possible for instance by considering (i) absolute sensing of the gradient, i.e. $\mathbf{v} = \chi \nabla a$, (ii) assuming an upper sensing threshold $K$ with Michaelis-Menten kinetics, i.e. $\mathbf{v} = \chi \nabla a/(a+K)$ [9], or (iii) considering a finite logarithmic sensing regime within lower and upper attractant concentrations $K_-$ and $K_+$, respectively, i.e. $\mathbf{v} = \chi \nabla \log[(1+a/K_-)/(1+a/K_+)]$ [6, 10, 11]. For small $K_-$ or small $K$, both the finite logarithmic sensing and the Michaelis-Menten sensing will converge to $\nabla a/a$. In general, if the drift velocity vanishes quickly as $a \to 0$, e.g. for absolute sensing, Michaelis-Menten kinetics with large $K$, or finite logarithmic sensing with large $K_{-}$, we find that the migration patterns become less chemotactic and traveling waves cannot be formed, see the discussion in Section 2 below.

As explained in the main text, we can nondimensionalize the Eqs.(S1-S2) by $t \to (m\bar{\rho}_c)^{-1}t'$ and $x \to \sqrt{\frac{D_a}{m\bar{\rho}_c}}x'$, with a reference cell density $\bar{\rho}_c$. This spatiotemporal rescaling then leads to the nondimensional coupled PDE system:

$$\partial_t \rho_i = \tilde{D}_i \nabla^2 \rho_i - \tilde{\chi}_i \nabla \cdot \left( \rho_i \frac{\nabla a}{a} \right), \tag{S3}$$

and

$$\partial_t a = \nabla^2 a - \rho_c a, \tag{S4}$$

where $\tilde{D}_i \equiv D_i/D_a$ and $\tilde{\chi}_i \equiv \chi_i/D_a$ are the rescaled diffusion and chemotactic coefficients.

#### 1.1 Analytical prediction for the traveling wave velocity

For the system with cell influx, we numerically find a travelling wave-like propagation of cell density profiles with a well-defined velocity. Here we derive a simple analytical expression to estimate the velocity of this travelling wave front. For coupled consumer and sensor cell populations, this velocity is controlled solely by the consumer cells that locally shape the attractant gradient. We can therefore aim to find this velocity by considering the front-like propagation of the consumer cells and the attractant dynamics. To seek for self-similar density profiles in time, we switch to a comoving frame $z \equiv x - \mathscr{V}t$ with the front speed $\mathscr{V}$, and rewrite Eqs.(S1-S2) as:

$$-\mathscr{V}\rho_c' = D_c \rho_c'' - \chi_c \left( \rho_c \frac{a'}{a} \right)' \tag{S5}$$

and

$$-\mathscr{V}a' = D_a a'' - m\rho_c a, \tag{S6}$$

where prime denotes $d/dz$. As the consumer density behind the leading front is constant "in the bulk", i.e. $\rho_c = \rho_c^\dagger = const.$, we make the ansatz $a \propto \exp(\lambda z)$ and obtain

$$-\mathscr{V}\lambda = D_a \lambda^2 - m\rho_c^\dagger. \tag{S7}$$

Integrating Eq.(S5) from $z^\dagger$ inside the bulk to $z = \infty$, subject to conditions $\rho_c(z^\dagger) = \rho_c^\dagger$, and $\rho_c(\infty) = \rho_c'(z^\dagger) = \rho_c'(\infty) = 0$, then leads to:

$$\lambda = \mathscr{V}/\chi_c. \tag{S8}$$

Note that the diffusion term vanishes due to the constant (i.e. "flat") bulk density. Using Eq.(S7) we thus obtain for the front speed

$$\mathscr{V} = \chi_c \sqrt{\frac{m\rho_c^\dagger}{D_a + \chi_c}}, \tag{S9}$$

or in the nondimensionalized system, leading to

$$\mathscr{V} = \tilde{\chi}_c \sqrt{\frac{\rho_c^\dagger}{1 + \tilde{\chi}_c}}. \tag{S10}$$

Interestingly, this result indicates that the Keller-Segel system can exhibit traveling wave solutions where the velocity is selected by the boundary influx, which fixes the bulk density $\rho_c^\dagger$. Therefore, the system does not rely on the Kolmogorov criterion to have sharply localized initial cell densities to select a speed, in contrast with Fisher-KPP waves [12]. From a biological perspective, this can be a robust mechanism to facilitate wave-like migration through the interaction with a chemical field. A similar expression for the traveling wave speed had been found in a model of angiogenesis [13] in the absence of cell influx, where the cell density at the left boundary ($z = 0$) was assumed to be fixed. We also note that an expression for the front speed was recently obtained for bacterial chemotaxis with cell growth [9], which did not explicitly depend on the time scale of attractant consumption given by $(m\rho_c^\dagger)^{-1}$, but on the growth rate of bacteria in the bulk, highlighting how cell influx vs. growth might complementarily act to generate traveling waves.

#### 1.2 Chemoattractant kinetics in closed vs. open systems

Time evolution of the chemoattractant density as governed by Eq.(S2) indicates that at steady state, attractant concentration becomes $a^{\text{st}} = 0$ as there is no external supply to balance its consumption by the chemotactic cell population. With respect to chemoattractant kinetics, this choice thus corresponds to a *closed* system with no in- or outflux of the attractant between the bulk of the system and the exterior. Experimentally, migration assays in a microfluidic channel with two holes at the ends approximate such a closed system, as after equilibration there are no external reservoirs to further supply chemoattractants into the migration channel. Alternatively, we can envisage an *open* system, where attractant molecules can enter or exit the system through the boundaries, leading to an effective turnover kinetics in addition to the consumption by cells. This choice thus corresponds experimentally to under-agarose migration assays, as the confined cell migration zone is in constant contact with a large reservoir of chemoattractant [14]. More generally, any setup where the cell migration zone is surrounded by semi-permeable boundaries that only allow the chemoattractant molecules to be transferred would be described by such turnover kinetics. In the context of immune cell migration in vivo, for instance, these two choices describe two limits where the chemoattractant molecules are either in fixed amount or constantly replenished along the path of migration.

In addition to the internalization by consumer-sensor cells, time evolution of the chemoattractant concentration in an open system then becomes:

$$\partial_t a = D_a \nabla^2 a + \underbrace{r - ka}_{\text{turnover}} - \underbrace{m\rho_c a}_{\text{uptake by cells}} \quad , \tag{S11}$$

where $r$ is a target concentration rate that describes the attractant influx into the system, and $k$ is an effective "decay" rate for the outflux or loss of chemoattractants.

**Nondimensionalization.** Depending on the choice of closed vs. open systems, i.e. in the absence vs. presence of a chemoattractant turnover term in Eq.(S11), we obtain different spatiotemporal rescaling factors for nondimensionalizing the coupled PDE system. For the open case with nonzero turnover in Eq.(S11), we can rescale the cell and attractant concentrations to reduce the independent parameters $r$ and $k$. Introducing the transformations $t \to k^{-1}t'$, $x \to \sqrt{\frac{D_a}{k}}x'$, $\rho_c \to \frac{k}{m}\rho_c'$ and $a \to \frac{r}{k}a'$, and after dropping the primes we obtain:

$$\partial_t \rho_i = \tilde{D}_i \nabla^2 \rho_i - \tilde{\chi}_i \nabla \cdot \left( \rho_i \frac{\nabla a}{a} \right), \tag{S12}$$

and

$$\partial_t a = \nabla^2 a + 1 - a - \rho_c a, \tag{S13}$$

where again $\tilde{D}_i \equiv D_i/D_a$ and $\tilde{\chi}_i \equiv \chi_i/D_a$ are the reduced control parameters, as in the closed case. Note that, even though we have the two additional parameters $r$ and $k$ for the chemoattractant turnover, the nondimensionalized form of the open system is still controlled by the four rescaled variables $\tilde{D}_i$ and $\tilde{\chi}_i$.

#### 2 Model predictions and parameter sensitivity

Here we discuss some features of the model for different choices of control parameters, and provide results on the transient dynamics of the system.

**Single-population dynamics.** The simplest example for chemotaxis via self-generated gradients can be explored by looking at the dynamics of a single consumer cell type. The nondimensional system of equations then indicate that the consumer migration is controlled by the rescaled diffusion and chemotactic coefficients $\tilde{D}_c$ and $\tilde{\chi}_c$, respectively. We find in particular that the ratio $\tilde{\chi}_c/\tilde{D}_c$ (or $\chi_c/D_c$) is the key control parameter that describes the transition of consumers from exhibiting diffusive-like to chemotactic migration profiles. Indeed, for $\chi_c/D_c > 0.4$, we found that the spatial profiles of consumers show a well-defined density peak, see Fig.S1A. To define such a peak, we use the convention that the cell density at the boundary should not be larger than half of the maximal cell density, i.e. $\rho(x=0) < \rho_{max}/2$. Furthermore, in this chemotactic regime, the long-time scaling of the mean position $\langle x \rangle \propto t^\alpha$ had exponents much larger than for simple diffusion, with typically $\alpha > 0.65$. In contrast, for $\chi_c/D_c < 0.4$ density peaks were not as pronounced and generically exhibited $\rho_{max}/2 < \rho(x=0)$, while the scaling exponent decayed to $\alpha \leq 0.65$ and converged to $\alpha = 0.5$ for $\chi_c/D_c \to 0$, for instance being equal to $\alpha \simeq 0.55$ for $\chi_c/D_c = 0.1$, see Fig.S1B.

**Variations in the diffusion coefficients.** Including the second population of sensor cells in the system, we first wanted to test the influence of variations in their random motility as controlled by the rescaled diffusion coefficient $\tilde{D}_s$. Setting $\chi_c/D_c = 3$ for sufficiently chemotactic consumer cells, and identical chemotactic coefficients for consumers and sensors e.g. $\tilde{\chi}_s = \tilde{\chi}_c$, we found that a large diffusion coefficient for sensors (e.g. $D_s/D_c = 5$) led to slowly spreading densities for the sensor cell population, with a scaling exponent of $\alpha_s < \alpha_c$, which resulted in sensor cells falling behind the consumer population at long times (see Fig.S1C). Furthermore, we found that decreasing the diffusion coefficient of the sensor cells to be smaller than that of the consumers ($D_s/D_c < 1$) did not influence the coupled vs. uncoupled regimes, where $\chi_s/\chi_c < 1$ led to mean position ratios

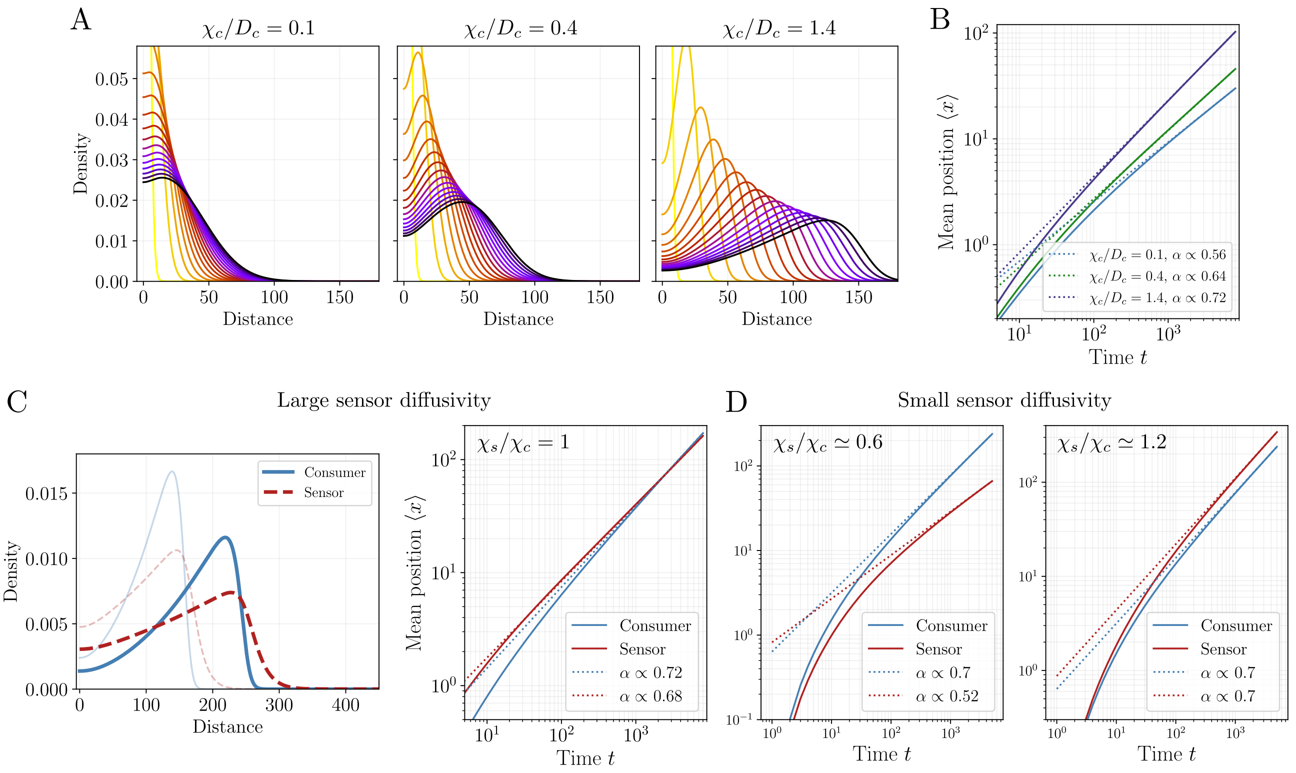

Supplementary Figure S1: **Model predictions for different parameter regimes.** A-B) Migration dynamics for a single consumer cell population. A) Spatial cell density profiles over time (time points color-coded) for three different choices of consumer chemotactic strength $\chi_c/D_c$. For $\chi_c/D_c \simeq 0.4$, the cell density on the left boundary remains relatively high, exceeding the half-peak density $\rho_{max}/2$ (left and middle panels), in contrast with the strongly chemotactic case with $\chi_c/D_c = 1.4$, where a well-defined density peak can be observed (right panel). B) Mean position of the cell population as a function of time ($\langle x \rangle \propto t^{\alpha}$) shows larger scaling exponents $\alpha$ with increasing $\chi_c/D_c$. C-D) Influence of sensor diffusion coefficient $D_s$ on the migration patterns. C) Cell density profiles (left) of the consumer and sensor population for identical chemotactic coefficients $\chi_c = \chi_s$ but with a large rescaled diffusion coefficient for the sensor population with $D_s/D_c = 5$. Long-time scaling of the mean position (right) shows that the sensor population has a smaller exponent $\alpha_s < \alpha_c$ and eventually falls behind the consumer cell population. D) Mean position scaling for the case when the sensor population has a smaller diffusion coeffcient than that of the consumers with $D_s/D_c = 0.083$. For $\chi_s/\chi_c \simeq 0.6$ (left), i.e. in the uncoupled regime, the sensor population falls behind the consumer cells with $\alpha_s < \alpha_c$. For $\chi_s/\chi_c \simeq 1.2$, i.e. in the coupled regime, sensors can propagate ahead of the consumer cell popoulation where the scaling exponents match with $\alpha_s = \alpha_c \simeq 0.7$.

of $\bar{x} \equiv \langle x_s \rangle / \langle x_c \rangle < 1$ with scaling exponents $\alpha_s < \alpha_c$ in the uncoupled regime, and for $\chi_s/\chi_c \geq 1$ we always recovered $\bar{x} > 1$ with $\alpha_s \simeq \alpha_c$ indicating the long-time coupling (see Fig.S1D). Finally, for identical consumer and sensor diffusibilities ($\tilde{D}_s = \tilde{D}_c$) the phase diagram of mean position ratios exhibited $\bar{x} < 1$ in the entire uncoupled region ($\chi_s/\chi_c < 1$) for all choices of consumer chemotactic ability $\chi_c/D_c$ (see Fig.S2A).

**Slow-time dynamics of the system.** Even though the migration dynamics in the absence of cell influx necessarily consists of slowly decaying density profiles, i.e. transient migration patterns, we found that the phase diagram as well as the scaling dynamics were preserved over long times, see Fig.S2C for a comparison of the mean position ratio phase diagram at different time points. Indeed, we found that the density profiles could be

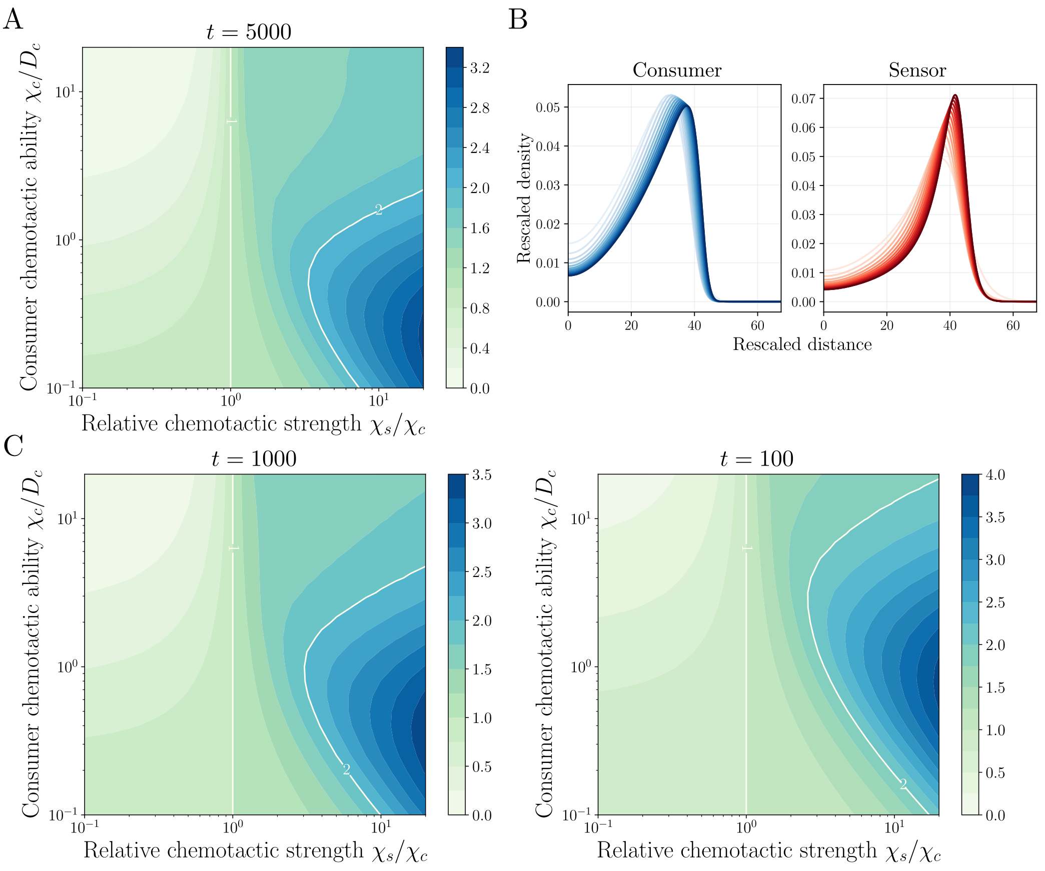

Supplementary Figure S2: **Temporal dynamics of the migration patterns.** A) Phase diagram of mean position ratios $\bar{x} \equiv \langle x_s \rangle / \langle x_c \rangle$ at a late time point ($t = 5000$) for the case of identical rescaled diffusion coefficients for sensor and consumer cell populations with $\tilde{D}_s = \tilde{D}_c = 0.1$. In contrast with the phase diagram shown in the main text (where $\tilde{D}_s \neq \tilde{D}_c$), for identical diffusion coefficients the mean position ratio fulfils $\bar{x} < 1$ in the entire uncoupled regime with $\chi_s/\chi_c < 1$. The remaining features of the phase diagram are largely preserved, in particular the bounded increase of $\bar{x}$ for sufficiently chemotactic consumer cells with a large chemotactic ability $\chi_c/D_c$. B) Density profiles of both consumer (left) and sensor (right) populations exhibit an approximately scale-invariant form over distinct times (shaded colors) using transformations $x \to x(t^{\varphi})$ and $\rho_i \to \rho_i(t^{\theta})$ with appropriate rescaling exponents $\varphi$ and $\theta$. C) Phase diagrams of mean position ratios $\bar{x}$ at early time points of $t = 1000$ (left) and $t = 100$ (right) exhibit qualitatively similar features as in the long-time limit, with most changes observed at early times for large $\chi_s/\chi_c$.

approximately mapped onto a scale-invariant form with appropriate rescaling factors, see Fig.S2B. This suggests that the diffusive leakage of cells from the propagating front does not influence the coupling mechanism over long times.

**Uncoupled regime with cell influx.** We next asked whether the predictions for the uncoupled regime with $\chi_s/\chi_c < 1$, where sensor cells fell behind the consumer cell population (as shown in Fig.1 in the main text), also held true for the system with nonzero influx of cells at the left boundary, i.e. $\partial_x \rho|_{x=0} = \gamma$. We used the same parameter set for consumers and sensors as in Fig.3 of the main text and only changed the sensor chemotactic coefficient to be slightly smaller than the consumer chemotactic strength with $\tilde{\chi}_s = 0.24$ and $\tilde{\chi}_c = 0.3$. This already led to a markedly diffferent migration pattern for the sensors: Spatial profiles did not exhibit any density

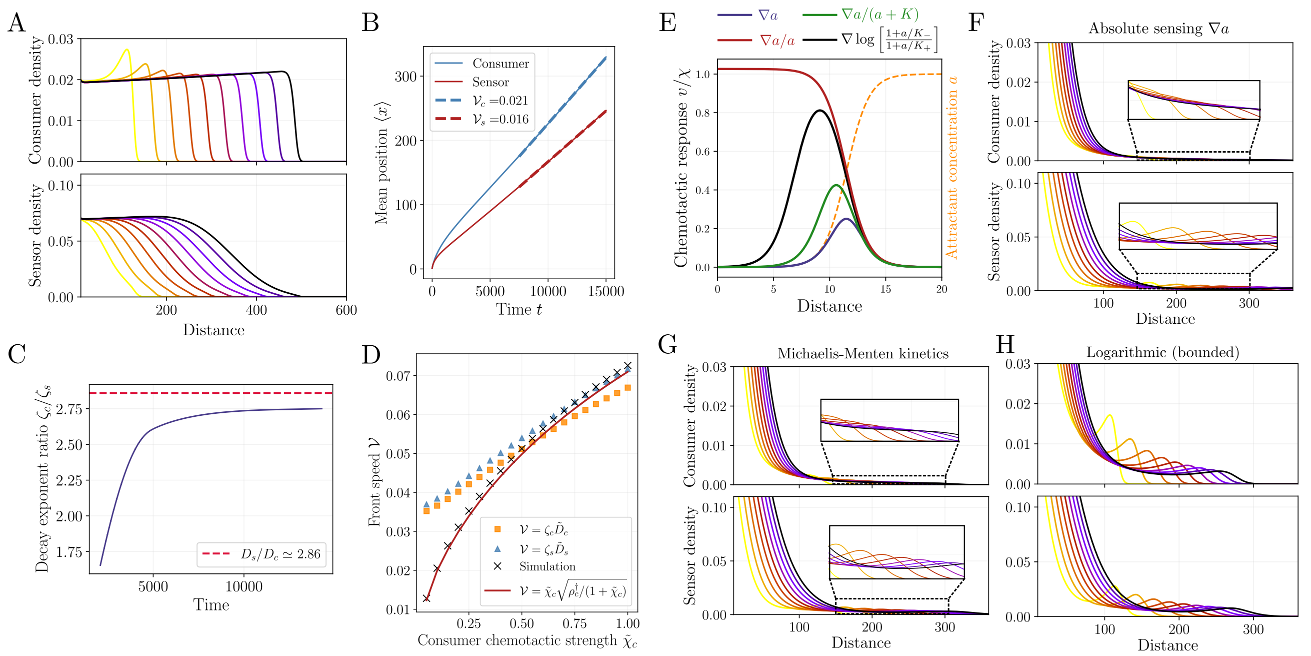

Supplementary Figure S3: **Dependence of travelling wave solutions on different parameter choices and sensing functions in the closed system.** A) Traveling wave profiles for different time points (color coded) in the case of an uncoupled migration pattern with $\chi_s < \chi_c$, where sensor cells lag behind the propagating consumer cell front (with $\tilde{\chi}_c = 0.3$ and $\tilde{\chi}_s = 0.24$). B) Mean position of cell densities over time indicate that the velocity-coupling breaks down for $\chi_s < \chi_c$. The consumer cell population propagates with a larger velocity ahead of the sensor cell population. C) Numerical test for estimating the diffusion coefficient ratio of the two cell populations $D_s/D_c$ from their decay profiles given by the relation $\rho_i \propto \exp(-\zeta_i z)$ with $\zeta_i = \mathscr{V}/\tilde{D}_i$. Decay exponents fitted for the two cell populations at every time point exhibit a ratio $\zeta_c/\zeta_s$ over time that approaches the diffusion coefficient ratio $D_s/D_c$. D) Numerical test for the analytical prediction of the traveling wave velocity $\mathscr{V}$ (given by Eq.(S10)) for different choices of the rescaled consumer chemotactic coefficient $\tilde{\chi}_c$. Analytical estimate (red line) very closely approximates the wave speeds obtained from the numerical solution of the PDEs (crosses). For comparison, velocity estimates inferred from the decay profiles of cell densities by the relation $\mathscr{V} = \zeta_i \tilde{D}_i$ (squre and triangular markers) fail to describe the numerical wave speed for small $\tilde{\chi}_c$. E) Illustration of different chemotactic response functions given by $v/\chi$, where $v \equiv |\mathbf{v}|$ is the drift velocity. For an attractant profile with an exponential tail (dashed line), absolute (purple) and relative (red) gradient sensing, Michaelis-Menten kinetics with half-maximum concentration $K$ (green), as well as bounded logarithmic sensing (black) within upper and lower ranges $K_+$ and $K_-$ are shown. F-H) Time evolution of density profiles for the cases of absolute sensing (F), Michaelis-Menten kinetics (G), and bounded logarithmic sensing (H). Only in the latter case, when the ratio $K_-/K_+$ is sufficiently small, we observe transient density peaks of consumer cells, where the cells in the 'bulk' are partially recruited to the leading front. Parameter choices for the different cases are: $\tilde{\chi}_c = 3$, $\tilde{\chi}_s = 3.6$ for (F) and (G); $\tilde{\chi}_c = 0.3$, $\tilde{\chi}_s = 0.36$ and $\tilde{K} \equiv K_-/K_+ = 0.01$ for (H), and the rescaled diffusion coefficients are $\tilde{D}_c = 0.07$ and $\tilde{D}_s = 0.2$ for all cases.

peaks at any time point and the majority of the sensor density remained in the back of the consumer front, see Fig.S3A. Furthermore, the mean position of cell densities over time indicated that the velocity coupling observed in the coupled regime disappeared in the uncoupled regime: Sensor cells propagated with a smaller velocity than the consumer cell population over long times, see Fig.S3B.

**Spatial decay exponents of the density profiles.** As we observed traveling waves for the case of cell influx through the boundary, we could switch to a comoving frame as a standard method to analyze stationary features [15], and arrive at the simple expression for the cell density given by Eq.(S5). At the leading tail of the traveling front, i.e. for saturated flat regions of the chemoattractant concentration with $a' \simeq 0$, this equation then dictates that the density profiles should scale as

$$\rho_i \propto \exp(-\zeta_i z) = \exp(-\mathscr{V} z/\tilde{D}_i)\,, \tag{S14}$$

where the subscript $i = c, s$ denotes the consumer or sensor population. This means that both cell densities will have exponential tails with decay lengths given by $\zeta_i \propto \tilde{D}_i^{-1}$. We can thus use this information to directly read off the relative diffusion coefficients from the density profiles of the cell populations using $\zeta_c/\zeta_s = D_s/D_c$. To test this prediction, we first looked at the scaling of cell density profiles around sufficiently flat regions of the chemoattractant concentration, i.e. for large $z$ values. We first confirmed that density profiles indeed exhibited long exponential tails at saturating regions of the attractant gradient (i.e. ahead of the propagating fronts), as qualitatively predicted from the theory. After determining the decay exponents $\zeta_i$ from exponential fits to the density profiles, we then calculated the ratio $\zeta_c/\zeta_s$ to see if this reproduced the ratio of diffusion coefficients used as input. From the numerical solution for the time evolution of density profiles we observed that the decay exponent ratio indeed approached $D_s/D_c$, see Fig.S3C.

**Numerical test of the traveling wave velocity.** A concrete prediction from the analytical result for the front velocity, as given in Eq.(S10), is that it should scale with the square-root of the consumer cell chemotactic coefficient, i.e. $\mathscr{V} \propto \sqrt{\tilde{\chi}_c}$. To test this prediction, we systematically varied the rescaled chemotactic coefficient $\tilde{\chi}_c$ of the consumer cells and numerically calculated the front speed. We then compared the numerical values with the analytical prediction given by Eq.(S10), where we determined the bulk consumer density $\rho_c^\dagger$ from the final shape of the numerical density profiles. We found that the analytical prediction very closely matched the numerical values for the front speed for the entire range of $\tilde{\chi}_c$ values, see Fig.S3D. We then checked to what degree the approximate relation $\mathscr{V} = \zeta_i D_i$, see Eq.(S14), which strictly applies only for small chemoattractant gradients ($a' \simeq 0$), held in this range of parameters. After determining the decay exponents $\zeta_i$ from exponential fits to the spatial density profiles, we then compared this estimate with the numerical and analytical values for the front speed, and found that it only provided a good approximation for sufficiently large chemotactic coefficients $\tilde{\chi}_c$ but failed to reproduce the correct speeds otherwise, see Fig.S3D.

**Influence of different chemotactic response functions on the travelling wave pattern.** We next turned to test the robustness of the traveling wave solutions to different types of chemotactic response functions, as the cells' sensing ability might be constrained by the absolute concentration of the chemoattractant. In the minimal logarithmic response defined by $\mathbf{v} = \chi \nabla \log(a)$, for instance, as $a \to 0$ the cells would still have a constant response, e.g. for an exponentially decaying attractant profile. Therefore, we now briefly discuss three additional response functions given by (i) absolute gradient sensing $\mathbf{v} = \chi \nabla a$, (ii) sensing dictated by

| Parameter | Estimate |
|---|---|
| Rescaled consumer diffusion coefficient $\tilde{D}_c$ | 0.07 |
| Rescaled sensor diffusion coefficient $\tilde{D}_s$ | 0.2 |
| Rescaled consumer chemotactic coefficient $\tilde{\chi}_c$ | $0.3^{\dagger}$, $1.5^{\ddagger}$ |
| Rescaled sensor chemotactic coefficient $\tilde{\chi}_s$ | $0.36^{\dagger}$, $2.25^{\ddagger}$ |
| Cell influx rate $\gamma$ | -0.01 |
| Temporal rescaling factor $\eta = (m\bar{\rho}_c)^{-1}$ | $0.2\text{min}^{\dagger}$ |
| Temporal rescaling factor $k^{-1}$ | $0.06\text{min}^{\ddagger}$ |

Table S1: **Parameter values used in the numerical evaluation of the nondimensional system of equations.** Dagger symbols ($^{\dagger}$) indicate parameter values used for the comparison with the data from microfluidic channel experiments, while double-dagger symbols ($^{\ddagger}$) correspond to estimates for the comparison with the under-agarose experiments. Other parameter values used in the analysis of alternative model predictions are indicated in the corresponding figure captions.

Michaelis-Menten kinetics $\mathbf{v} = \chi \nabla a/(a+K)$ with half-maximum concentration $K$, and (iii) logarithmic sensing $\mathbf{v} = \chi \nabla \log[(1+a/K_-)/(1+a/K_+)]$ bounded within lower and upper concentration limits $K_-$ and $K_+$, respectively. Fig.S3E illustrates the different response functions for an attractant profile with an exponentially decaying tail given by $a \propto \exp(x)/(C+\exp(x))$, where $C$ is a sufficiently large constant.

We then asked whether these different forms of chemotactic response functions had an influence on the migration patterns, in particular with respect to the existence and dynamics of traveling wave profiles. To explore these cases, we first nondimensionalized the coupled PDEs for each choice of chemotactic sensing function. This could be done by introducing a rescaling for the attractant concentration as $a \to Ka'$ for the Michaelis-Menten kinetics, and as $a \to K_+ a'$ for bounded logarithmic sensing. The corresponding nondimensional equations for the cell density evolution then followed $\partial_t \rho_i = \tilde{D}_i \nabla^2 \rho_i - \tilde{\chi}_i \nabla \cdot (\rho_i \nabla a/(a+1))$ and $\partial_t \rho_i = \tilde{D}_i \nabla^2 \rho_i - \tilde{\chi}_i \nabla \cdot (\rho_i \nabla \log[(1+a/\tilde{K})/(1+a)])$, respectively. The latter equation for logarithmic sensing thus introduces a new rescaled parameter $\tilde{K} = K_-/K_+$ that is given by the ratio of the lower and upper concentrations.

Numerical solutions of the PDEs showed that deviations from the relative sensing led to notable changes in the cell density evolution: First of all, we did not observe traveling wave solutions for any of the alternative sensing mechanisms. Next, both absolute sensing and Michaelis-Menten kinetics resulted in accumulation of cells close to the boundary of the system, and even for the case of highly chemotactic cell populations with $\tilde{\chi}_c/\tilde{D}_c \simeq 40$, consumer cells did not form well-defined density peaks, see Fig.S3F-G (insets). In contrast, bounded logarithmic sensing with a small $K_-/K_+$ ratio led to transient peaks of consumer cells that decayed slowly in time, see Fig.S3H, indicating that in this case cells in the bulk could be partially recruited to the leading front.

### 3 Parameter estimates and perturbation experiments

In this section we briefly outline the inference of key parameters of the system, and describe perturbation experiments performed to test the predictions of the theory. Parameter estimates that are used for the comparison with experimental data are summarized in Table S1.

#### 3.1 Inference of diffusion coefficients

As the nondimensionalized system is entirely controlled by the rescaled diffusion and chemotactic coefficients $\tilde{D} \equiv D/D_a$ and $\tilde{\chi} \equiv \chi/D_a$, where $D_a$ denotes the diffusion coefficient of the attractant, estimating the diffusion coefficients of the consumer and sensor cell populations is a key step to constrain the analysis for the experimentally relevant migration regime. We approached this using complementary methods to obtain reproducible estimates for the diffusion coefficients.

**Diffusion coefficient of consumer/dendritic cells.** (i) First, we looked at the migration data of dendritic cells (DCs) in under-agarose experiments in the absence of the chemoattractant CCL19, as published previously [14]. From the scaling of mean-squared deviations (MSD) of each cell trajectory $\langle (r(T) - r(0))^2 \rangle = 4D_cT$, we first analyzed trajectories that had reached $T = 120$ mins to focus on sufficiently processive cells, and estimated diffusion coefficient values from 2 different experiments with a mean of approximately $D_c \simeq 390 \pm 55 \mu m^2/\text{min}$ ($\pm$ standard error (SE)) (see Fig.S4A). (ii) Second, we determined the scaling of MSD of all trajectories (from $N = 173$ cell trajectories) over time (up to $T = 120$ mins and considering trajectories that were processive enough to exceed $\sim 50\mu$m radial distance), and used Furth's formula for persistent random walks [16] to fit this dataset:

$$\langle (r(t) - r(0))^2 \rangle = 2nD_c(t - \tau(1 - e^{-t/\tau})) \,, \tag{S15}$$

where $n = 2$ is the dimension of the system and $\tau$ the persistence time of cells. We found that the average MSD data could be well-fitted with $D_c \simeq 300\mu m^2/\text{min}$ and a persistence time of $\tau \simeq 30$ mins (see Fig.S4B). (iii) Third, we turned to the velocity autocorrelation function (averaged over $N = 173$ trajectories) and calculated the diffusion coefficient using

$$nD_c = \int_0^T dt \langle \mathbf{v}(t) \cdot \mathbf{v}(0) \rangle \,, \tag{S16}$$

which, after integrating over $T = 120$ mins, gave us an estimate of $D_c \simeq 292\mu m^2/\text{min}$ (see Fig.S4C). These three methods therefore indicated a consistent range for the diffusion coefficient of the DCs. As the MSD scaling given by Furth's formula involves fitting of two parameters, we assumed that it presumably slightly underestimated the diffusion coefficient. Therefore, we decided to take $D_c \simeq 350\mu m^2/\text{min}$ to account for the upper limit, which was also closer to values reported previously for dendritic cells [17]. Using a chemoattractant diffusion coefficient of $D_a \simeq 86\mu m^2/s$ [14], we could then fix the rescaled diffusion coefficient of DCs by $\tilde{D}_c = D_c/D_a \simeq 0.07$.

**Diffusion coefficient of sensor / T cells.** To determine the diffusion coefficient of the T cells, we used experimental datasets from under-agarose experiments where DCs and T cells migrated in a uniform field of the

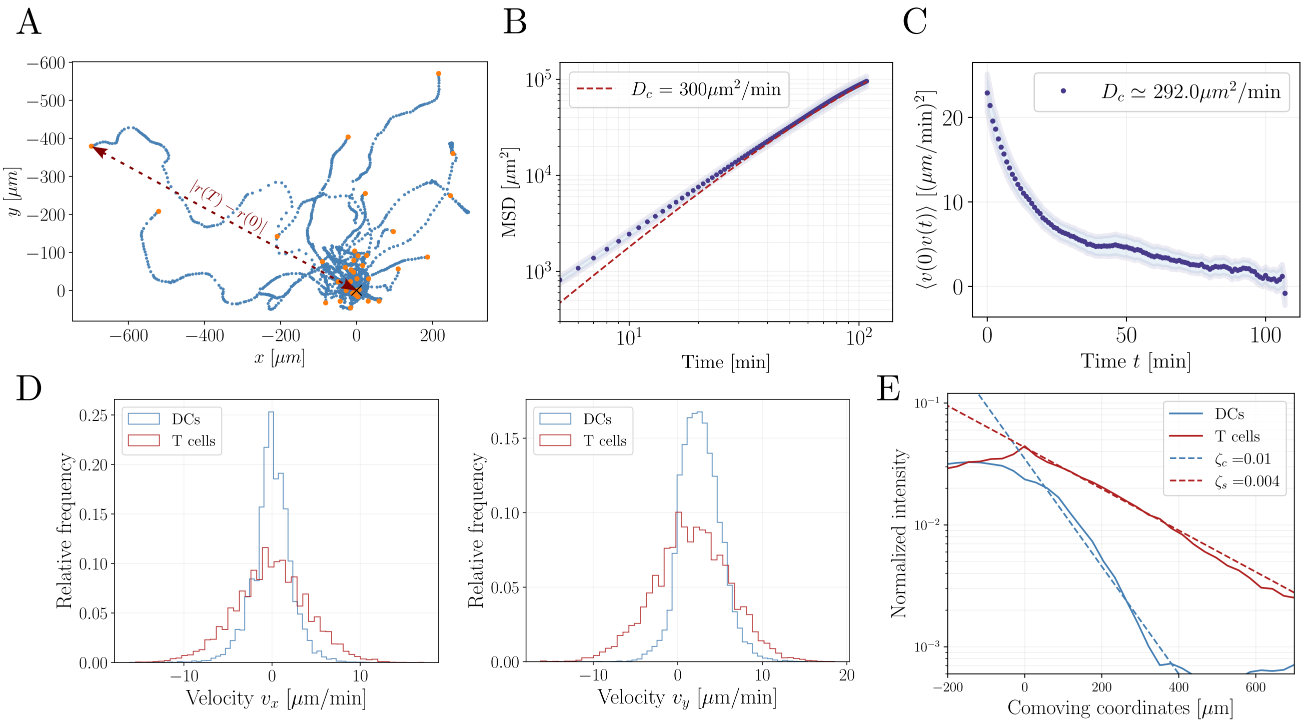

Supplementary Figure S4: **Estimation of the rescaled diffusion coefficients of DC (consumer) and T cell (sensor) populations.** A) Spider plot of DC trajectories from under-agarose experiments in the absence of the chemoattractant CCL19. For each cell trajectory, orange dots indicate the position of the corresponding cell at time point $T = 120$min, whereas blue dots describe its past coordinates. The MSD is then calculated from the absolute distance $|r(T) - r(0)|$ between its final $\mathbf{r}(T)$ and initial $\mathbf{r}(0)$ coordinates, which led to an estimate of $D_c \simeq 390\mu m^2/$min. B) Alternative calculation of the MSD by taking into consideration of each time point in a given cell trajectory. For each time interval $\Delta t$, MSD is calculated using a "sliding window" for each cell trajectory. Average MSD datasets (circles) obtained from $N = 173$ cell trajectories are plotted over increasing time intervals and compared against a fit using Eq.(S16) (dashed red line). C) Average velocity autocorrelation function against time from $N = 173$ cell trajectories. The estimated diffusion coefficient from the integral of the velocity correlation function ($D_c \simeq 292\mu m^2/$min) is close to the fitted value ($D_c \simeq 300\mu m^2/$min) in (B). Error bars in (B-C) represent SEs. D) Histogram of dendritic (blue) and T cell (red) velocities obtained from the under-agarose assays in the presence of the chemoattractant CCL19. Both the $x-$ (left) and $y-$ (right) components of velocities show comparable fluctuations for each cell population, whereas T cell velocities have notably larger SDs than that of the DCs with $\sigma(v_s) \simeq 4.3\mu m/min$ vs. $\sigma(v_c) \simeq 2.3\mu m/min$, indicating a diffusion coefficient ratio of $D_s/D_c = \sigma(v_s)^2/\sigma(v_c)^2 \simeq 3.5$. E) Semi-log plot of the average density profiles of DCs (blue) and T cells (red) in the comoving frame of the T cell peak, obtained from the microfluidic channel experiments (profiles correspond to the one displayed in Fig.3D in the main text). Decaying tails of the profiles ($\rho_i \propto \exp(-\zeta_i z)$) are fitted by the decay exponents $\zeta_c = 0.01$ and $\zeta_s = 0.004$, which indicate a diffusion coefficient ratio of $D_s/D_c \simeq 2.5$. Experimental data plotted in (A-D) from [14].

chemoattractant CCL19, as published in [14]. As the T cell trajectories in this setup can potentially involve interactions with the DCs particularly in dense regions, we decided not to use the MSD or velocity autocorrelation data to estimate the T cell diffusion coefficient. Instead, we looked at the distributions of velocity components in the two ($x$ & $y$) directions and compared their fluctuations both for the DC and T cell populations, see Fig.S4D for the velocity distributions. As the diffusion coefficient of the cells is proportional to the variance of velocity distributions, regardless of advective fluxes driven by the coupling to the chemoattractant, we could then infer the relative diffusion coefficient of the T cells (sensors) via:

$$D_s/D_c \propto \frac{\sigma(v_s)^2}{\sigma(v_c)^2} \equiv \frac{\langle (v_s - \langle v_s \rangle)^2 \rangle}{\langle (v_c - \langle v_c \rangle)^2 \rangle}\,. \tag{S17}$$

Interestingly, even though the mean velocities in the $y$ direction reflect the bias due to advective chemotactic flux, we could separate the diffusive contribution as the variances in $v_x$ and $v_y$ exhibit similar values: The variance in $v_x$ distributions were $\sigma(v_{s,x})^2 \simeq 17(\mu m/\text{min})^2$ for T cells and $\sigma(v_{c,x})^2 \simeq 5(\mu m/\text{min})^2$ for DCs, and the variance of the $v_y$ distributions were $\sigma(v_{s,y})^2 \simeq 20(\mu m/\text{min})^2$ for T cells and $\sigma(v_{c,y})^2 \simeq 6(\mu m/\text{min})^2$ for DCs. This analysis indicated that the ratio between the diffusion coefficients of the sensor (T cell) and consumer (DC) populations was about $D_s/D_c \simeq 3.5$.

Furthermore, we considered another independent method to infer the ratio of diffusion coefficients for the case of travelling waves from the decay profiles of cell densities, as predicted from Eq.(S14) and numerically tested in Fig.S3C. Because traveling waves are obtained in the microfluidic channel experiments, we could then simply fit the decay lengths of the density profiles in the comoving frame with the travelling wave velocity $\mathscr{V}$. We found that the ratio of the decay lengths between the T cell (sensor) and DC (consumer) populations fell consistently within the range $\zeta_c/\zeta_s = D_s/D_c \simeq 2.5$ (se Fig.S4E), in good agreement with the diffusion ratio obtained from the variance of velocity distributions from the under-agarose experiments. These two independent methods from different experimental setups led us to conclude that the diffusion coefficient ratio of sensor and consumer populations is approximately given by $D_s/D_c \simeq 3$. Using the rescaled diffusion coefficient for consumer cells of $\tilde{D}_c = 0.07$, as estimated above, we then used the value $\tilde{D}_s = 0.2$ for the rescaled diffusion coefficient of the sensor cells.

#### 3.2 Parameter fitting for the comparison between theory and experiment

After estimating the rescaled diffusion coefficients $\tilde{D}_i$, the dynamics of the nondimensional system of equations is entirely controlled by the rescaled chemotactic coefficients $\tilde{\chi}_i$ together with the choice of initial and boundary conditions. As stated in the main text, we first numerically observed a traveling wave solution when there is a nonzero boundary flux of cells at $x = 0$. We used a small influx rate $\gamma = 0.01$ for comparison with both experimental setups, although this rate can be changed if a more quantitative match between absolute density values is needed. As we did not have quantitative data on the exact densities from the microfluidic channel experiments, but used relative intensity datasets instead, we did not tune this influx rate further.

We observed for the closed system that if the relative chemotactic strength between sensor and consumer cells was $\chi_s/\chi_c > 2$, sensor cells accumulated ahead of the consumers in ever increasing numbers over time. This constrained the chemotactic ratio to be within $\chi_s/\chi_c \simeq 1-2$. The choice for the rescaled chemotactic coefficient of the consumer cell population was relatively unconstrained, as any sufficiently large value with $\chi_c/D_c > 2$ led to traveling waves in the closed system. We therefore used $\chi_c/D_c \simeq 4$ to obtain a well-defined wave. In contrast, for the open system with attractant turnover, we chose $\chi_c/D_c \simeq 20$ to reproduce the experimental density profiles, although the relative chemotactic strength of the two cell populations was chosen to be $\chi_s/\chi_c \simeq 1.5$, similar to the closed system case. The initial conditions for the attractant profile in both systems were fixed by a constant spatial density with $a(x,t=0)=1$. For the consumer and sensor cell populations we used sharply localized initial density profiles with exponentially decaying tails described by $\rho_i(x,t=0)=1/(1+A\exp(x-B))$ with $A=B=1$ for the closed system, and $A=1$ and $B=5$ for the open system.

Finally, to transform the nondimensional solutions to physical spatiotemporal units, we had to fix the rescaling factors in $t \to \eta t'$ for the closed system and in $t \to k^{-1}t'$ for the open system. To reproduce the experimentally observed migration dynamics we then set $\eta = 0.2$min and $k^{-1} = 0.06$min, which then also rescaled the spatial dimensions via $x \to \sqrt{D_a\eta}x'$ and $x \to \sqrt{D_a k^{-1}}x'$ for the closed and open systems, respectively. Table S1 summarizes the parameter estimates used for the comparison between numerical solutions and experiments.

##### 3.3 Co-migration of dendritic cells and CCR7-KO T cells

One key prediction of the theory is that the coupled migration of the consumer and sensor cell populations is strictly controlled by their relative chemotactic coefficients, i.e. by the ratio $\chi_s/\chi_c$. More precisely, we showed that the coupled regime breaks down for $\chi_s/\chi_c < 1$ where the sensor population cannot form a well-defined peak ahead of the consumer front and falls behind. To test this prediction experimentally, we decided to use CCR7-KO T cells, which lack the receptor CCR7 that binds the chemoattractant CCL19, mixed with WT DCs to see whether their migration efficiency would be influenced by attractant-specific responses. Using microfluidic channel experiments we found that CCR7-KO T cells were not able to migrate ahead of the DCs and fell behind the DC front, even though they initially had a mean position ahead of DCs, see Fig.S5A-C and Supplementary Movie 2. Their velocity was also notably smaller than that of the DC front, indicating the breakdown of the velocity-coupling of traveling waves we observed in the WT experiments. Interestingly, the DCs migrated again in an approximately well-preserved density profile with a constant velocity of $5.3\mu m/min$, Fig.S5D, close to their traveling wave velocity when they migrated with WT T cells. This suggests that T cells do not act as a mechanical drag on the DC motility, and excludes such mechanical interactions between the DCs and T cells as the regulatory mechanism behind the coupled migration pattern we observed in the WT experiments.

##### 3.4 T cell migration in uniform and imposed gradients

Another simple prediction of the model is that, because the sensors / T cells cannot shape their own guidance signals, they should not be able to migrate as efficiently alone, i.e. in the absence of a consumer / DC population. In fact, concentration peaks of sensor populations should only exist in the mixed system with a gradient-modulating cell population. To test these predictions, we looked at the migration patterns of only T cells in microfluidic channel experiments both in uniform (equilibrated) as well as gradient setups for the chemoattractant CCL19. First, we could not observe any well-defined density peaks in either uniform or gradient setups. Second, we found that although the T cells showed quite rapid motion in a uniform CCL19 field as in the mixed setup (see Supplementary Movie 3), this was mainly due to their random, undirected movements. Indeed, their mean velocity was

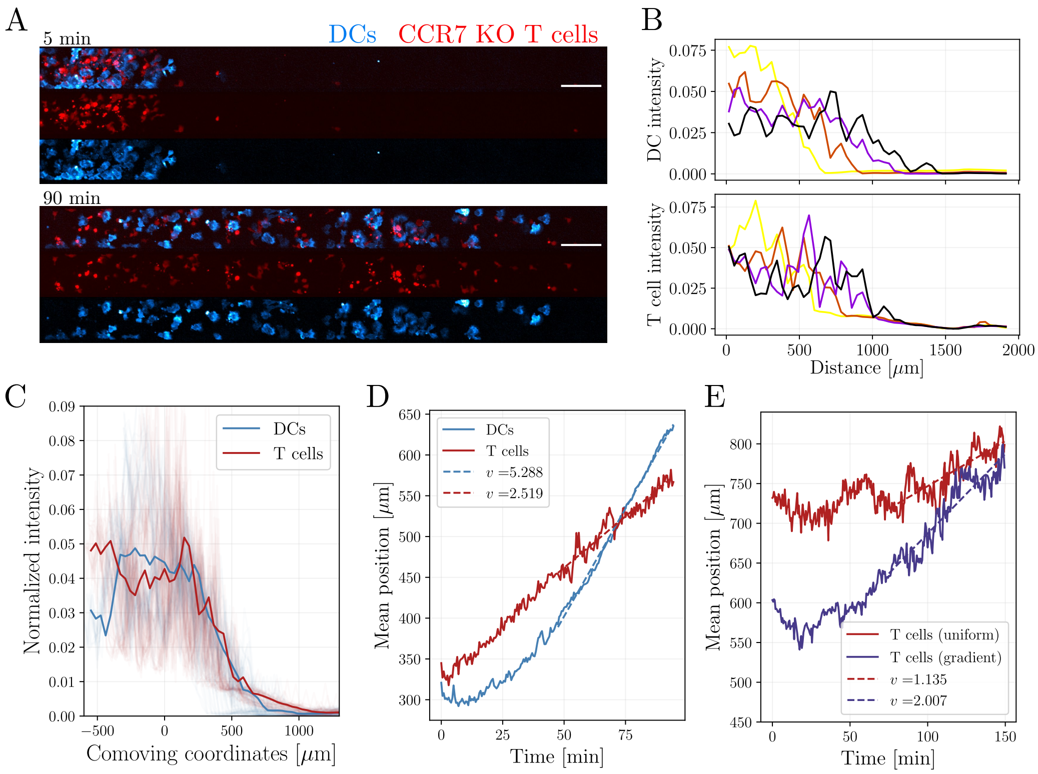

Supplementary Figure S5: **Perturbation experiments to test coupled migration dynamics of cell populations.** A) Microscopy images from the microfluidic channel experiments with labelled DCs (blue) and co-migrating CCR7-KO T cells (red) at $t = 5$min (top) and $t = 90$min (bottom). Scale bar indicates $100\mu$m. B) Averaged normalized intensities for DCs (top) and CCR7-KO T cells (bottom) from $n = 2$ experiments at different time points (color-coded). C) Intensity profiles at different time points overlaid in the refence frame co-moving with the mean DC position. Mean density profile of CCR7-KO T cells (red) does not exhibit a well-defined peak, while most T cell population being placed behind the propagating DC front, in contrast with the WT case shown in Fig.3 of the main text. D) Mean position of cell densities over time for DCs (blue) and CCR7-KO T cells (red) show the breakdown of the velocity coupling, where T cells now fall behind the DC population and migrate with a smaller velocity at long times. E) Mean position of WT T cell populations in the absence of a co-migrating DC population, obtained from microfluidic channel experiments with an initially uniformly distributed CCL19 field (red) and a gradient of CCL19 (purple). In both cases, the average velocity obtained from the mean position is notably smaller than that of the WT T cell populations co-migrating with DCs.

about $1\mu m/min$ in the uniform field (see Fig.S5E, red line), strongly deviating from their dynamics in the mixed setup where they co-migrated with DCs with velocities of $5-6\mu m/min$. Interestingly, even in the pre-patterned gradient case, their mean velocity remained around $2\mu m/min$ (see Fig.S5E, purple line), suggesting that the velocity-coupling in the mixed setup provides an efficient way for the sensors to migrate over long distances.

#### 4 Migration patterns in the open system

In the open system in contact with an external chemoattractant reservoir, additional to its local internalization by consumer cells, the attractant evolution is regulated by a turnover rate that shifts its global level to a target concentration, see Eq.(S11). Therefore, the local gradients dynamically shaped by the consumers will be directly

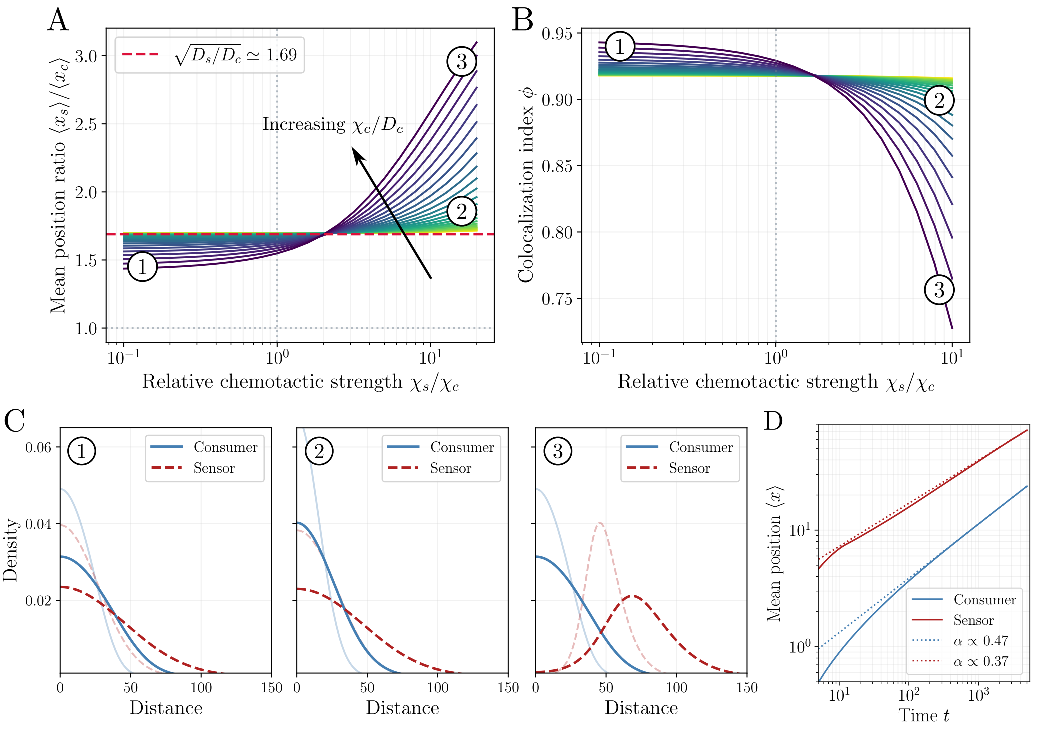

Supplementary Figure S6: **Migration and colocalization patterns in the open system with nonzero attractant turnover.** A) Mean position ratio $x \equiv= \langle x_s\rangle/\langle x_c\rangle$ as a function of relative chemotactic strength $\chi_s/\chi_c$ between sensor and cconsumer populations for different values of chemotaxis-to-diffusion ratio of consumers (color-coded). In the uncoupled regime with $\chi_s < \chi_c$, mean position ratio is largely independent of $\chi_c/D_c$ and takes values close to the that dictated by diffusive spreading, i.e. $\bar{x} \simeq \sqrt{D_s/D_c}$. For strongly chemotactic consumers with a large $\chi_c/D_c$, increasing the chemotactic strength of sensors (large $\chi_s/\chi_c$) leads to deviations from diffusive spreading with $\bar{x} > \sqrt{D_s/D_c}$. B) Colocalization between consumer and sensor population as quantified by the index $\phi$ is maximal in the uncoupled regime with $\chi_s < \chi_c$ and monotonically decays with increasing $\chi_s/\chi_c$. For strongly chemotactic consumer cells with a large $\chi_c/D_c$, the colocalization is most sensitive to variations in the relative chemotactic strength $\chi_s/\chi_c$. C) Cell density profiles from different parameter regions plotted in (A) and (B) exhibit the emergence of pulse-like propagation for sensor cells only for large $\chi_c/D_c$ and large $\chi_s/\chi_c$. Consumer cells exhibit similar profiles for all parameter choices, and do not form pulse-like patterns. Parameter values used for the density profiles were $\tilde{\chi}_c \simeq 0.014$ and $\tilde{\chi}_s \simeq 0.28$ (Region 1); $\tilde{\chi}_c \simeq 1.4$ and $\tilde{\chi}_s \simeq 0.18$ (Region 2); $\tilde{\chi}_c \simeq 1.4$ and $\tilde{\chi}_s \simeq 22$ (Region 3). D) For strongly chemotactic consumer and sensor populations with $\chi_c/D_c \simeq 20$ and $\chi_s/\chi_c \simeq 16$, mean position of cell densities over time ($\bar{x} \propto t^{\alpha}$) shows that the long-time scaling exponents are below 0.5 and $\alpha_s < \alpha_c$.

influenced by these additional effects. To explore this, we numerically evaluated the nondimensional Eq.(S12) and analyzed the migration patterns in different parameter regions.

**Phase diagram of relative positions and colocalization.** We first looked at how the sensor-to-consumer mean position ratio $\bar{x} \equiv \langle x_s\rangle/\langle x_c\rangle$ was influenced in the open system: We found that in the uncoupled regime with $\chi_s < \chi_c$, where sensor cells are less chemotactic than consumers, mean position ratio always attained values close to $\bar{x} \simeq \sqrt{D_s/D_c}$ dictated by diffusion regardless of the consumer chemotactic strength $\chi_c/D_c$ (see Fig.S6A). This is in strong contrast with the behavior we observed for the closed system, where increasing $\chi_c/D_c$ in the uncoupled regime leads to a strong decrease in $\bar{x}$, because consumer cells can form a well-defined density peak and migrate arbitrarily ahead of sensors. For large $\chi_s/\chi_c$ and strongly chemotactic consumer cells with large $\chi_c/D_c$ in the open system, we observed that sensor cells now could propagate ahead of consumer in a "pulse-like" density (see Fig.S6C), leading to mean position ratios $\bar{x} > \sqrt{D_s/D_c}$ (see Fig.S6A). Furthermore, density profiles over time showed that consumer cells never formed density peaks and were spreading over time with a large concentration confined at the boundary, see Fig.S6C for the spatiotemporal profiles at different parameter regimes.

Next, we analyzed the colocalization patterns of the two cell populations by using the colocalization metric $\phi$ based on Jensen-Shannon divergence (see Section 5 below for details). We found that in the uncoupled regime with $\chi_s < \chi_c$ the colocalization was maximal with values $\phi > 0.9$, indicating that this regime was indeed dominated by the diffusive behavior of the two cell populations (see Fig.S6B and density profiles in Region 1 in Fig.S6C). Large values of the consumer chemotactic strength $\chi_c/D_c$ only resulted in negligible changes in $\phi$. For large $\chi_c/D_c$, increasing the relative chemotactic strength of sensors, i.e. for large $\chi_s/\chi_c$, led to a strong decrease in colocalization due to the pulse-like propagation of the sensors ahead of the consumer cells (see density profiles in Regions 2 and 3 in Fig.S6C).

To better understand the dynamics of the open system where a sensor cell pulse could be formed, we looked at the mean position of the two cell populations over time for large $\chi_c/D_c$ and large $\chi_s/\chi_c$. We found that despite the relatively large value chosen for the chemotactic strength of consumers ($\chi_c/D_c \simeq 20$), the scaling exponent for the mean position (via $\langle x_c \rangle \propto t^{\alpha_c}$) exhibited values around $\alpha_c \simeq 0.47$, indicating slightly sub-diffusive migration, see Fig.S6D. This was in strong contrast with the closed system, where $\chi_c/D_c \simeq 20$ leads to scaling exponents of $\alpha \simeq 0.7$. Furthermore, the scaling exponent of the sensor cells was around $\alpha_s \simeq 0.37 < \alpha_c$, which implied that sensors were being slowed down by the (sub-)diffusive behavior of the consumers in the back of the sensor peak.

#### 5 Jensen-Shannon divergence for colocalization

In addition to the co-migration efficiency of coupled cell populations, another important question is to what degree these different populations spatially overlap or interact with each other, as this would be the precondition for any mechanical or contact-based signaling interactions. Particularly in the context of immune response, frequent physical contacts between DCs and T cells allow DCs to present antigens and to activate T cells [18]. To quantify the spatial overlap between cell populations, we reasoned that the colocalization metric should encode both the spatial proximity as well as the similarity in the shapes of the density profiles. For instance, the spatial overlap between a pulse-like and diffusive density profile should indicate a smaller colocalization than the overlap between two pulse-like or two diffusive profiles, as the latter two cases would lead to maximal overlap of densities. Such a metric can be defined by considering the cell concentrations as probability distributions $\rho_i(x) \to P_i(x)$ with a suitable normalization, i.e. $P_i(x) \equiv \rho_i(x)/\sum_x \rho_i(x)$, where $i = c, s$ denotes the consumer and sensor cell populations, respectively. We can then use the Jensen-Shannon divergence [19] to describe the

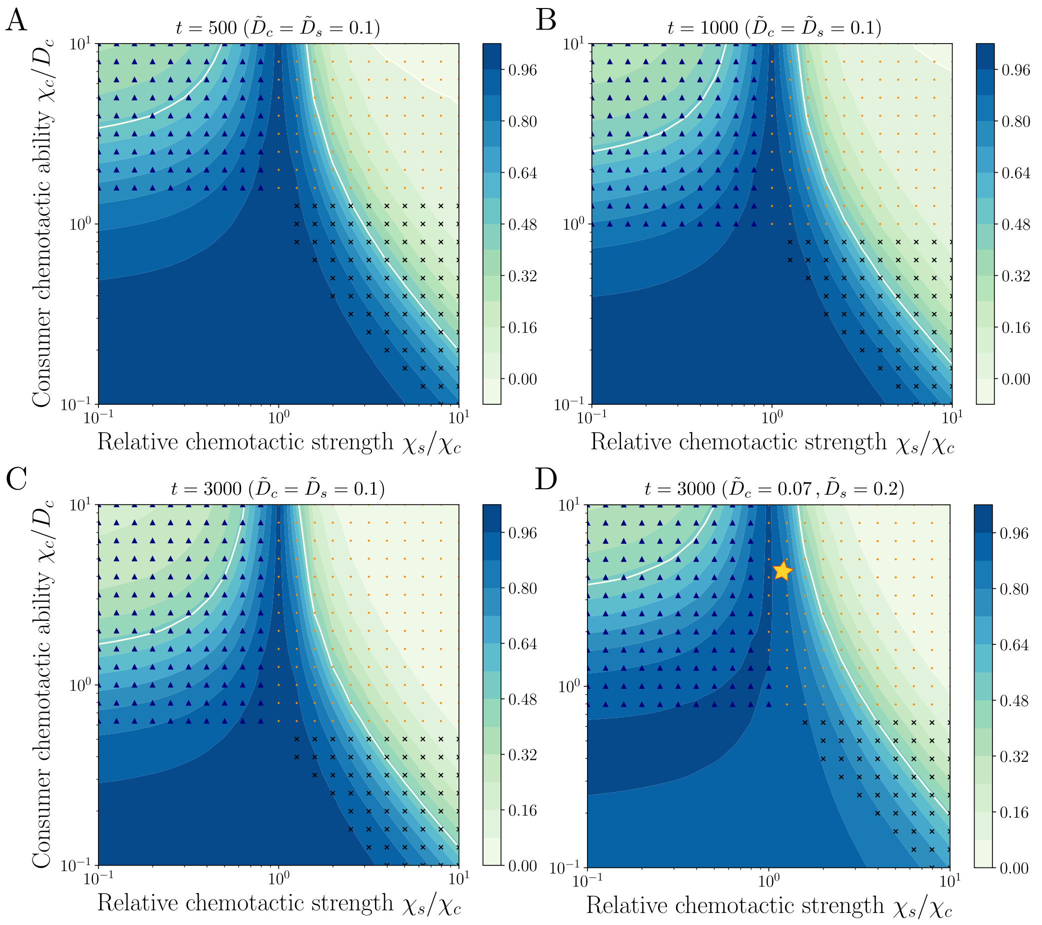

Supplementary Figure S7: **Stability of the phase diagram for the colocalization of consumer and sensor cell populations.** A-C) Colocalization index $\phi$ evaluated from the cell density profiles at time points $t = 500$ (A), $t = 1000$ (B), and $t = 3000$ (C) for the case of identical diffusion coefficients $\tilde{D}_c = \tilde{D}_s = 0.1$. The phase diagram remains largely preserved in particular in the coupled regime with $\chi_s/\chi_c > 1$. As $t$ increases, density profile peaks for consumer (triangular markers) and both cell types (dots) occur in larger regions of the parameter space. D) For different diffusion coefficients as inferred from the experimental data (with $\tilde{D}_c = 0.07$ and $\tilde{D}_s = 0.2$), the colocalization is mainly influenced in the uncoupled regime of the phase space with $\chi_s/\chi_c < 1$. For weakly chemotactic consumer cells with $\chi_c/D_c < 1$ the colocalization of both cell populations is reduced in comparison with the case of identical diffusion coefficients. The parameter set for the comparison with the experimental system (star symbol) indicates a colocalization of $\phi \simeq 0.9$.

similarity between the two probability distributions, defined by

$$D_{JS}(P_c, P_s) \equiv \frac{1}{2}\left(D_{KL}(P_c, M) + D_{KL}(P_s, M)\right), \tag{S18}$$

where $M(x) = \frac{1}{2}\left(P_c(x) + P_s(x)\right)$ and $D_{KL}(P,Q) \equiv \sum_x P(x) \log\left(\frac{P(x)}{Q(x)}\right)$ is the Kullback-Leibler divergence. Note that, we do not make use of divergence metrics for continuous random variables, as we discretize space in finite intervals of $\Delta x$ to obtain the solutions of the coupled PDE system (see below for details on the numerical method). Finally, as $D_{JS} = 0$ indicates maximal similarity while $D_{JS} = 1$ corresponds to zero overlap (using base 2 logarithm), we use the more intuitive choice $\phi = 1 - D_{JS}$ as a metric for colocalization.

Because the comparison of cell densities requires fixing a certain time point $t = t'$, their colocalization $\phi$ will in principle change dynamically as the cells migrate over space. However, as we had found for the phase diagram

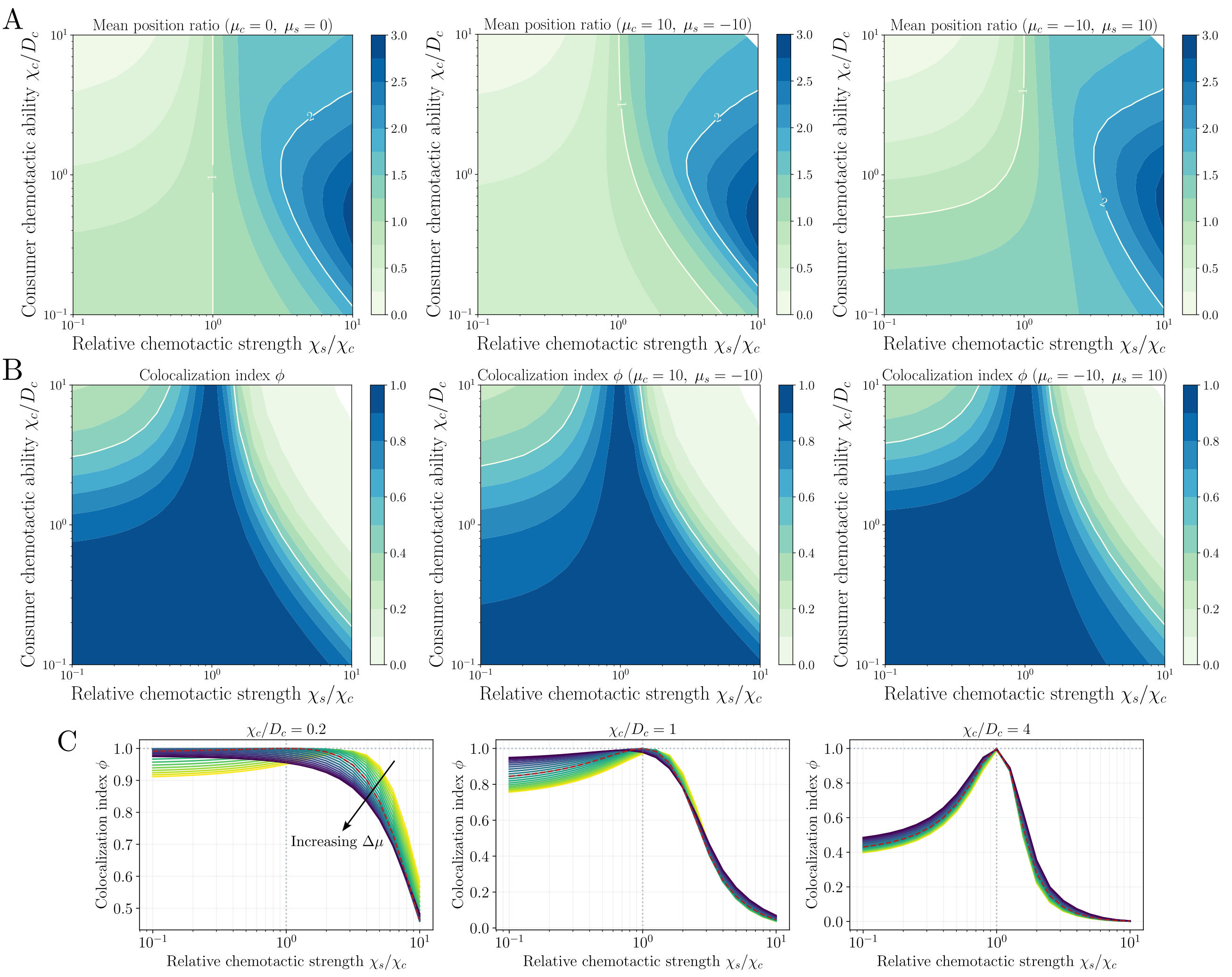

Supplementary Figure S8: **Influence of non-reciprocal mechanical interactions on the mean position ratio and colocalization of cell populations.** (A) Mean position ratio of cell populations for different choices of non-reciprocal mechanical coupling $\Delta\mu = \mu_s - \mu_c$. Phase diagram of mean positions changes primarily for weakly chemotactic consumer cells with $\chi_c/D_c \leq 1$ in the presence of mechanical interactions. Contour line delineating equal mean position ratio at $\chi_s/\chi_c = 1$ without mechanical coupling ($\Delta\mu = 0$, left panel) shifts to larger $\chi_s/\chi_c$ values for $\Delta\mu < 0$ (middle panel) and to smaller $\chi_s/\chi_c$ values for $\Delta\mu > 0$ (right panel) in for weak consumer chemotaxis. (B) Colocalization index $\phi$ phase diagram shows small changes for small $\Delta\mu < 0$ (middle panel) and large $\Delta\mu > 0$ (right panel) in the weak consumer chemotaxis regime with $\chi_c/D_c \leq 1$. (C) Colocalization index for different values of the differential mechanical coupling parameter $\Delta\mu$ indicates that variations in $\Delta\mu$ only lead to small changes in the colocalization index $\phi$ for $\chi_c/D_c \leq 1$ (left and middle panales). Dashed line (red) represents the case $\Delta\mu = 0$. For strong consumer chemotaxis ($\chi_c/D_c = 4$, right panel) colocalization index converges to the predicted values in the absence of mechanical interactions with $\Delta\mu = 0$.

of relative mean positions, see Fig.S2, the phase space for colocalization $\phi$ also remains relatively preserved over time. Figs.S7A-C show the colocalization phase diagram evaluated at different time points for consumer and sensor cell populations with identical diffusion coefficients $\tilde{D}_c = \tilde{D}_s = 0.1$. As mentioned in the main text, using experimentally inferred values for the rescaled diffusion coefficients mainly shifts the colocalization values $\phi$ in the diffusive and uncoupled regime with $\chi_c/D_c < 1$ and $\chi_s/\chi_c < 1$, see Fig.S7D.

### 6 Mechanical interactions between cell populations

To explore the potential role of mechanical interactions such as cell-cell adhesion or density sensing between cell populations, we modified the coarse-grained chemotaxis description to include an additional advective flux term. We chose a minimal form that allows each cell population to linearly read off the density gradient of the other, and to become either attracted or repelled by it. In the nondimensionalized form, the equations for the consumer and sensor cell density evolution then read:

$$\begin{aligned} \partial_t \rho_c &= \tilde{D}_c \nabla^2 \rho_c - \tilde{\chi}_c \nabla \cdot (\rho_c \nabla a/a) + \tilde{\mu}_c \nabla \cdot (\rho_c \nabla \rho_s) \,, \\ \partial_t \rho_s &= \tilde{D}_s \nabla^2 \rho_s - \chi_s \nabla \cdot (\rho_s \nabla a/a) + \tilde{\mu}_s \nabla \cdot (\rho_s \nabla \rho_c) \,, \end{aligned} \tag{S19}$$

where $\tilde{\mu}_c \equiv \bar{\rho}_s \mu_c / D_a$ and $\tilde{\mu}_s \equiv \bar{\rho}_c \mu_s / D_a$ are the rescaled mechanical coupling parameters with the reference consumer and sensor cell densities $\bar{\rho}_c$ and $\bar{\rho}_s$, respectively. The rescalings for $t$, $x$, $D_i$ and $\chi_i$ can be performed analogously to the original system of equations without mechanical coupling (see Eqs.S3-S4). Note that, for clarity we will drop the tildes in the following (as is done in the main text). With this formulation, positive values ($\mu > 0$) indicate repulsion, and negative values ($\mu < 0$) indicate attraction by the other cell density. Even though this model simplifies the impact of more generic density- or contact-dependent mechanisms in collective cell migration [20], it allows for an effective screening to test the relative roles of mechanical and chemotactic interactions on the co-migration patterns. We could then numerically evaluate the system of equations Eq.(S19) together with Eq.(S4) and explore the influence of the mechanical coupling parameters $\mu_i$ on the migration patterns, see below for the details on initialization and the numerical approach.

To start testing the effect of mechanical interactions, we first set the diffusion coefficients of consumer and sensor cell types to be equal, i.e. $D_c = D_s = 0.1$, and looked at the mean position ratio $\langle x_s \rangle / \langle x_c \rangle$ of the two cell populations for different choices of the coupling parameters $\mu_i$. We mapped the phase diagram for the mean position ratio for mechanical coupling parameters that reflected a non-reciprocal coupling between the cell populations. For this, we defined the difference between the mechanical coupling parameters $\Delta\mu \equiv \mu_s - \mu_c$ while keeping $|\mu_c| = |\mu_s|$. We explored small $\Delta\mu < 0$ (large $\mu_c > 0$ and small $\mu_s < 0$), indicating consumers being pushed by sensors while the latter are attracted, and large $\Delta\mu > 0$ (with small $\mu_c < 0$ and large $\mu_s > 0$), which indicated consumers are pulled and sensors are pushed. We found that for both extreme choices, the mean position ratio was primarily influenced by mechanical interactions for weakly chemotactic consumer cell populations with $\chi_c / D_c < 1$, see Fig.S8A.

Next, we asked whether the colocalization between cell populations was sensitive to variations in the mechanical interactions, as these interactions directly influence the shape of cell density profiles. Surprisingly, for the same set of of $\Delta\mu$ values as before, we found that the colocalization index was not markedly influenced by mechanical interactions, even in the weakly chemotactic regime of consumer cells, see Fig.S8B. A finer parameter scan for different values of $\Delta\mu$ while changing the strength of consumer cell chemotaxis indicated that only for small $\chi_c / D_c \leq 1$, changes in $\Delta\mu$ led to minor variations in the colocalization index $\phi$, while for strong consumer chemotaxis with $\chi_c / D_c = 4$ mechanical coupling did not influence the colocalization index and converge to values predicted in the original system without mechanical interactions, see Fig.S8C.

#### 7 Details on numerical solution of the PDEs

**Numerical methods for chemotaxis.** To solve the coupled nondimensional PDE system in 1D (as given by Eqs.(S3-S4) and (S12-S13)), we use the finite difference method to approximate the spatial and temporal derivatives. In particular, we discretize spatial coordinates within boundaries $0 \leq x \leq L$ in $N+1$ intervals of equal size $\Delta x$ such that $x_n = n\Delta x$, with $n = 0,\ldots,N$. Similarly, we discretize time in intervals of size $\Delta t$ such that $t_k = k\Delta t$. For the time derivatives, we use the forward difference approximation such that $\partial_t \rho(x,t) \simeq (\rho(x_n,t_k+\Delta t) - \rho(x_n,t_k))/\Delta t$. We furthermore use von Neumann boundary conditions with $\partial_x \rho(x = 0,t) = \gamma_0$ and $\partial_x \rho(x = L,t) = \gamma_L$, where $\gamma_{0/L}$ define the boundary fluxes into the system. In general, we set $\gamma = 0$ (for both $x = 0$ and $x = L$) while for the case of cell influx leading to the travelling wave profiles, we choose a nonzero influx $\gamma$ at $x = 0$. As evaluating the second derivative at $x_0 = 0$ requires a value for $x^* = -\Delta x$ in the centered difference method, we introduce an additional lattice point for $n = -1$ and evaluate $\partial_x^2 \rho(0,t) \simeq 2(\rho(x_1,t_k) - \rho(x_0,t_k) - \gamma\Delta x)/\Delta x^2$ as an approximation at the boundary $x_0 = 0$. Likewise, the right boundary condition then dictates $\partial_x^2 \rho(N,t) \simeq 2(\rho(x_{N-1},t_k) - \rho(x_N,t_k) + \eta\Delta x)/\Delta x^2$, where $\partial_x \rho(x = L,t) = \gamma_L$. In all cases considered here, we will assume that there is no influx at the right boundary such that $\gamma_L = 0$.

We next initialize the consumer & sensor cell densities, as well as the chemoattractant concentration at $t = 0$. As we focus on chemotaxis by self-generated attractant gradients, we set the initial chemoattractant concentration to be uniformly distributed over space with a constant value, which we take for simplicity to be $a(x,t = 0) = 1$. We then initialize the consumer and cell density profiles to have sharply decaying profiles localized at $x = 0$, with initial values determined by $\rho_i(x,t = 0) = 1/(1 + A\exp(x - B))$, where $A$ and $B$ can be tuned to control the decay strength of the initial density profiles. Here, we generally use $A = B = 1$, except for the comparison with the experiments for the open system choose $A = 1$ and $B = 5$. We note that different choices with sharply localized and strongly decaying profiles in general lead to similar migration dynamics. After fixing the initial conditions we recursively update the cell and attractant profiles as determined by the finite difference approximations until a sufficiently large time $T \propto 10^3 - 10^4$ is reached. For stability we set the time interval to be $\Delta t = 0.01$ and the spatial discretization is set to $\Delta x = 1$.

**Numerical methods for chemotaxis with mechanical interactions.** To solve the PDE system corresponding to the case including mechanical interactions (see Eqs.(S19)), we first realized that if additional advective fluxes controlled by the mechanical parameter $\mu$ were too large, this led to numerical instabilities with the finite difference scheme for initially steep density profiles as chosen before. We therefore decided to use the buil-in function *NDSolve* in *Wolfram Mathematica 14.0* (Wolfram Research, Inc., Mathematica, Version 14.1, Champaign, IL (2024)). We furthermore chose smoother initial cell density profiles given by a Gaussian distribution $\rho_i(x,t = 0) = \sqrt{2\pi\sigma^2}^{-1} \exp(-\frac{1}{2}x^2/\sigma^2)$ with $\sigma = 10$. We first controlled whether this choice led to any changes in the dynamic evolution of the migration behavior in the absence of mechanical interactions and found that the phase diagrams remained largely unchanged, compare e.g. Fig.S2A with Fig.S8A (left panel), as well as Fig.4 (main text) with Fig.S8B (left panel). Having established numerical consistency, we then numerically solved the system of PDEs for different choices of mechanical coupling parameters $\mu_c$ and $\mu_s$ until a sufficiently large time of $T \propto 5000 - 10000$ is reached.

### 8 List of Supplementary Movies

- Supplementary Movie 1 - Collective Migration of DCs and T cells
- Supplementary Movie 2 - Collective Migration of DCs and CCR7 KO T cells
- Supplementary Movie 3 - T cell migration in uniform CCL19
